# DNA-binding mechanism and evolution of Replication Protein A

**DOI:** 10.1101/2022.07.20.500673

**Authors:** Clément Madru, Markel Martinez-Carranza, Sébastien Laurent, Alessandra C. Alberti, Maelenn Chevreuil, Bertrand Raynal, Ahmed Haouz, Rémy A. Le Meur, Marc Delarue, Didier Flament, Mart Krupovic, Pierre Legrand, Ludovic Sauguet

**Author notes:** To whom correspondence should be addressed (LS).

## Abstract

Replication Protein A (RPA) is a heterotrimeric single stranded DNA-binding protein with essential roles in DNA replication, recombination and repair, in both eukaryotic and archaeal cells. By using an integrative approach that combines three crystal structures, four cryo-EM structures in complex with single-stranded DNA (ssDNA) of different lengths, we extensively characterized RPA from *Pyrococcus abyssi* in different states. These structures show two essential features conserved in eukaryotes: a trimeric core and a module that promotes cooperative binding to ssDNA, as well as a newly identified archaeal-specific domain. These structures reveal for the first time how ssDNA is handed over from one RPA complex to the other, and uncover an unanticipated mechanism of self-association on ssDNA tracts. This work constitutes a significant step forward in the molecular understanding of the structure and DNA-binding mechanism of RPA, with far-reaching implications for the evolution of this primordial replication factor in Archaea and Eukarya.

## INTRODUCTION

In all forms of life, single-stranded DNA-binding proteins (SSBs) are essential components of the DNA replication machinery (Bain et al., 2018; Oliveira, 2021; Taib et al., 2021). They play vital roles in nearly all aspects of DNA metabolism by protecting exposed single-stranded DNA (ssDNA) and acting as platforms onto which DNA-processing enzymes can assemble (Brosey et al., 2013; Marceau, 2012). In Bacteria, SSB is the major single-stranded DNA binding protein. The archetypal *Escherichia coli* SSB encompasses a single oligonucleotide/oligosaccharide binding (OB) domain that assembles into homotetrameric complexes (Bianco, 2017). In Eukarya, the single-stranded DNA binding function is primarily achieved by the heterotrimeric Replication Protein A (RPA) complex (Wold, 1997). While RPA and SSBs share similar OB-fold DNA-binding domains, the heterotrimeric architecture of RPA is more complex than the homo-oligomeric assemblies of SSBs. Composed of three protein subunits, denoted as Rpa1, Rpa2, and Rpa3 (or RPA70, RPA32, and RPA14 in humans), RPA contains multiple OB-folds with different DNA-binding properties (**Figure 1a**). Over the past two decades, structural and biochemical studies have mostly been focused on the eukaryotic RPA by using X-ray crystallography (Bochkareva, 2002; Deng et al., 2007; Fan and Pavletich, 2012; Feldkamp et al., 2013; Jiang et al., 2006; Seeber et al., 2016), NMR (Brosey et al., 2009; Park, 2005), SAXS (Brosey et al., 2013), and cryo-electron microscopy (Yates et al., 2018).

**Figure 1:**
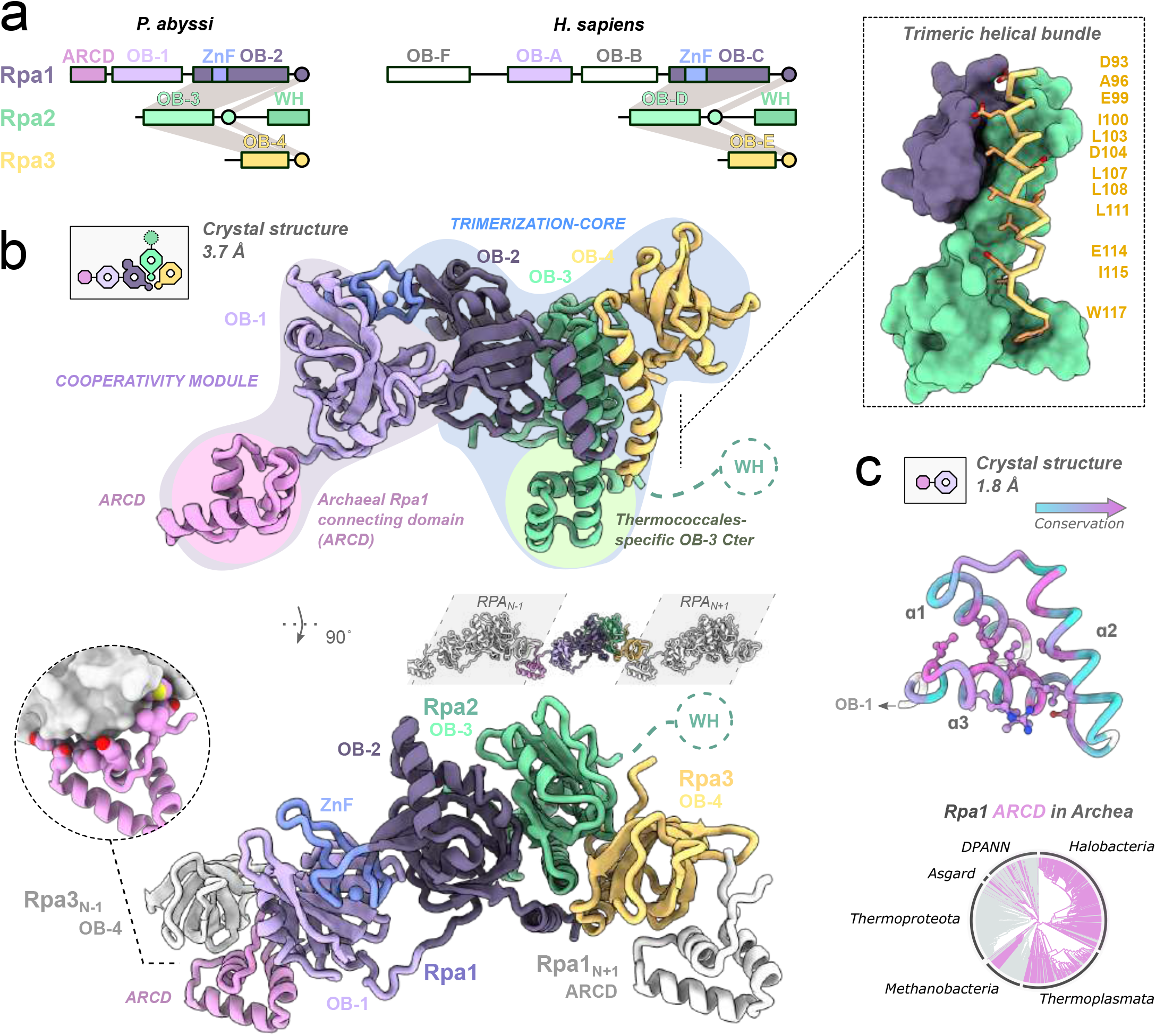
Structure of the archaeal RPA. **a**. Domain diagrams of PabRPA and human RPA. **b**. Two orthogonal views of the full-length PabRPA crystal structure at 3.7 Å with a focused view on the trimeric helical bundle (top right). Crystal lattice contacts within neighboring RPA heterotrimers are shown in grey. **c**. High resolution crystal structure of the Rpa1 cooperativity module at 1.8 Å (top) with a focused view on the newly identified archaeal-specific ARCD domain, which is colored according to sequence conservation. Side chains of residues connecting ARCD to OB-4 are shown as ball-and-sticks. Phylogenetic analysis (bottom) reveals a broad distribution of Rpa1-ARCD domain homologs across Archaea. Genomes lacking ARCD domain are presented in grey.

Archaea, which remains the most unexplored branch of the tree of life, possess an intriguing blend of bacterial and eukaryotic features as well as aspects that are unique to this domain of life (Duggin and Bell, 2006). While archaeal chromosomes resemble those of most Bacteria, their DNA replication machineries are closely related to their eukaryotic counterparts, serving as powerful models for understanding the function and evolution of the complex eukaryotic replication machineries. Thus, it is of considerable interest to understand how the simple bacterial-like chromosomes of Archaea are replicated by a eukaryotic-type replication apparatus (Barry and Bell, 2006). Interestingly, Archaea display a highly differentiated distribution of RPA, with a wide range of domain architectures with varying numbers of OB domains (MacNeill, 2021; Taib et al., 2021). No structure of archaeal RPA has been determined so far, except for structures of isolated OB domains that have been determined in the framework of structural genomics projects (PDBids: 2K50 and 3DM3). Determining the structure of a full-length archaeal RPA is required to fully appreciate its cellular function in the third domain of life.

We report three crystal structures and four cryo-EM structures of *P. abyssi* RPA (hereafter referred to PabRPA), in its apo form as well as bound to ssDNA substrates of different lengths. The structures include a trimeric core that is shared with eukaryotes and additional domains that are specific to Archaea. These structures reveal for the first time how ssDNA is handed over from one RPA molecule to the other. PabRPA is composed of two distinct modules, a trimerization core (Tri-C) with high-affinity for ssDNA and a second module located in the Rpa1 N-terminal region, which promotes cooperative binding to ssDNA. By using an integrative approach that combines X-ray crystallography, cryo-electron microscopy, and extensive biophysical analysis, we investigated the role of each individual domain of PabRPA and uncovered an unanticipated mechanism of self-association on ssDNA tracts, which has far-reaching implications for the DNA-binding mechanism of RPA in archaea and eukaryotes. We found that PabRPA forms tetrameric supercomplexes in the absence of DNA. In the presence of DNA, these tetramers undergo large conformational changes to form functional nucleoprotein filaments, which coat and protect ssDNA. This work also clarifies the evolutionary history of RPA, suggesting that RPA evolved in Archaea from a bacterial-like SSB ancestor, and that the eukaryotic RPA was inherited from an archaeal RPA ancestor.

## RESULTS

### Architecture of the archaeal RPA heterotrimeric complex

Due to its modular nature, RPA is extremely flexible and can adopt multiple conformations. X-ray crystallography studies have therefore focused on individual domains or truncated RPA trimeric cores (Bochkareva, 2002; Deng et al., 2007; Fan and Pavletich, 2012; Feldkamp et al., 2013; Jiang et al., 2006; Seeber et al., 2016). Here, we have biochemically reconstituted, crystallized and determined the 3.7 Å X-ray crystal structure of a heterotrimeric complex of *P. abyssi* Rpa1, Rpa2, and Rpa3 subunits (**Figure 1b**). An initial, partial model was obtained by using phase information derived from anomalous scattering data collected at the Zn K-edge. Model building was facilitated by determining two additional X-ray crystal structures of the N-terminal region of Rpa1 (1-180) at 1.8 Å resolution (**Figure 1c**) and the trimerization core bound to ssDNA at 3.2 Å resolution (**Supplementary figure S1 & Supplementary table S1**). Most of Rpa1(2-358), Rpa2(1-184) and Rpa3(6-117) subunits have been modelled in the electron density map, the only flexible region being the C-terminus of Rpa2(185-268). Bioinformatic studies have shown that this region contains a Winged-Helix (WH) domain that is conserved from Archaea to Eukarya, and recruits protein factors involved in DNA metabolism (MacNeill, 2021; Makarova and Koonin, 2013).

PabRPA contains four OB domains, named OB-1, OB-2, OB-3 and OB-4, which share the highest structural similarity with OB-A, OB-C, OB-D and OB-E in eukaryotic RPA, respectively (**Figure 1a**). Like in its eukaryotic OB-C counterparts, the amino acid sequence of PabRPA OB-2 contains a Zn-finger motif that is inserted between β-strands β1 and β2, and a short helical domain that is inserted between β-strands β3 and β4 (**Figure 1b & Supplementary figure S2**). The trimerization core (Tri-C) adopts a compact quaternary structure consisting of OB-2 of Rpa1, OB-3 of Rpa2, and OB-4 of Rpa3, which is the smallest subunit. Similar to the eukaryotic RPA, heterotrimerization of PabRPA is primarily mediated through a three-helix bundle formed by a C-terminal α-helix from each subunit (**Supplementary Movie S1**). PabRPA also displays intriguing specificities. Interestingly, the trimerization helix of Rpa2 is followed by a helix-turn-helix motif, acting as a pedestal that supports the trimerization helical bundle and stabilizes it by extending the contacts between the Rpa2 and Rpa3 subunits (**Figure 1b & Supplementary figure S2**). Thus, the total buried surface is 1209 Å^2^ for the PabRpa3 trimerization helix, but only 774 Å^2^ for its eukaryotic counterpart (PDBid: 4GOP). The chemical nature of the interactions is varied and includes a network of salt bridges and polar bonds, as well as extensive contacts between hydrophobic residues. Interestingly, the C-terminal extension of Rpa2 seems to be specific to Thermococcales, suggesting that it may play a role in further stabilizing the RPA trimerization core in organisms living at extremely high temperatures.

The N-terminal region of Rpa1 adopts an extended conformation that contrasts with the compact structure of the trimerization core and is shown here to play a critical role in the cooperative binding of PabRPA to ssDNA. Therefore, we named this independent DNA-binding module, the “cooperativity module”. The OB-1 domain of Rpa1 is bordered by an N-terminal tri-helical bundle, which has never been described so far and is referenced in the Pfam database (Mistry et al., 2021) as a domain of unknown function (DUF2240). This domain is absent in eukaryotes, but is broadly present in Archaea, being located at the N-terminal end of Rpa1 and Rpa1-like subunits, with very rare exceptions (**Figure 1c**). It is present in nearly all Thermoplasmata, Halobacteria and Asgard, but absent from Thermoproteota, which do not encode Rpa or Rpa-like proteins (**Supplementary figure S3**). In the following, we refer to this N-terminal domain as the Archaeal Rpa1 Connecting Domain (ARCD). Interestingly, in the PabRPA crystal structure, ARCD stacks against the OB-4 domain of neighboring Rpa3 chain within a lattice contact (**Figure 1b & Supplementary movie S1**). In the following sections, we describe the biological function of this new domain and show that it contributes to cooperative binding, by pre-connecting together RPA molecules before they bind to DNA.

### DNA-binding properties of PabRPA trimerization core and cooperativity module

The DNA-binding properties of PabRPA were investigated by electrophoretic mobility shift assays (EMSA), surface plasmon resonance (SPR) and biolayer interferometry (BLI) (**Figure 2a & Supplementary figure S4**). As expected, we found that PabRPA specifically binds to a 24-mer or a 32-mer random ssDNA with a dissociation constant in the low nanomolar range, but hardly binds to dsDNA or ssRNA (**Supplementary figure S4a-c**). As the use of homopolymeric ssDNA substrates is more convenient for structural studies, we verified that PabRPA binds to a poly-dT ssDNA, with an affinity similar to random ssDNA (1.25±0.6 nM vs 0.71±0.01 nM) (**Figure 2a & Supplementary figure S4d**). The DNA-binding properties of the PabRPA trimerization core and cooperativity module were further investigated by using BLI. Overall, our data show that binding to ssDNA is primarily ensured by the trimerization core. Yet, while both modules are required for an optimal binding to ssDNA, the trimerization core binds ssDNA with a 35-fold higher affinity (Kd=18.8±6.0nM) than the cooperativity module (Kd=650±150nM). Furthermore, the cooperativity module shows much faster dissociation kinetics than the trimerization core does. Indeed, about 60% of the PabRPA cooperativity module dissociates from ssDNA in 10 seconds, while in the same period, the trimerization core remains almost fully associated to ssDNA.

**Figure 2:**
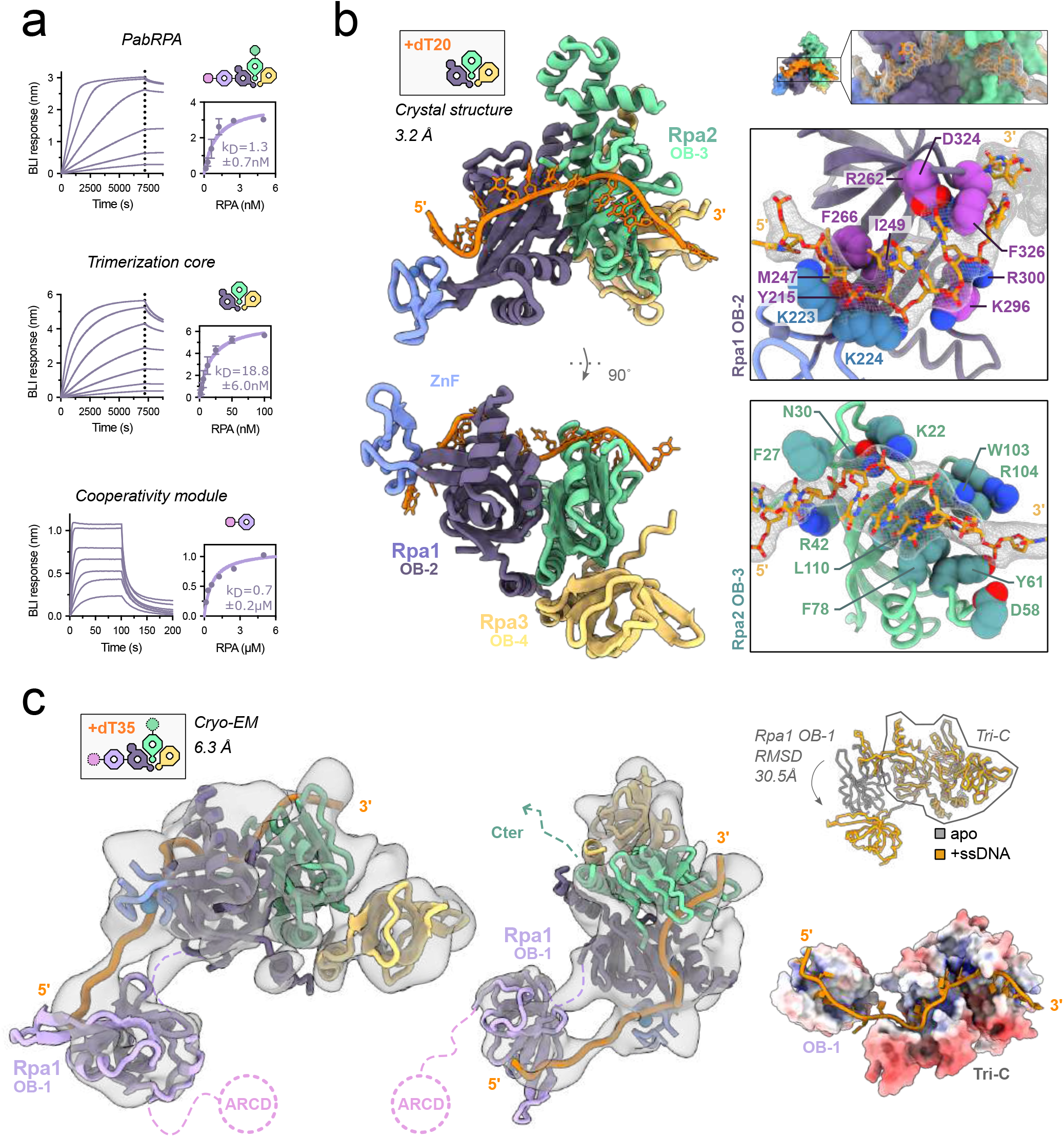
Structural basis for the DNA-binding activity by PabRPA. **a**. DNA-binding properties of PabRPA trimerization core and cooperativity module. Specific binding of PabRPA constructs to immobilized poly-dT35 ssDNA was measured by biolayer interferometry. Steady-state analysis were performed using the average signal measured at the end of the association steps. **b**. 3.2 Å crystal structure of the poly-dT20-bound Tri-C, with two focused views on Rpa1 OB-2 and Rpa2 OB-3 ssDNA-interacting residues. **c**. Cryo-EM structure of full-length PabRPA bound to a poly-dT35 ssDNA. Backbone traces of the poly-dT20-bound Tri-C and the Rpa1 OB-1 crystal structures were rigid-body fitted into the cryo-EM map contoured at a level of 1rmsd. Conformational changes of OB-1, observed between the apo-PabRPA crystal structure (in grey), and the DNA-bound cryo-EM structure (in orange). Electrostatic potential surface analysis showing that the repositioning of Rpa1 OB-1 domain extends the ssDNA-binding channel.

### Crystal structure of the PabRPA trimerization core bound to ssDNA

To gain insights into the structural basis for the high-affinity interaction between PabRPA and ssDNA, we determined the 3.2 Å crystal structure of the trimerization core bound to poly-dT20 ssDNA (**Figure 2b**). The modelled ssDNA lies in a channel that extends from the subunits Rpa1 to Rpa2, showing 14 contiguous nucleotides that traverse the OB-2 and OB-3 DNA-binding grooves. The DNA-binding grooves are lined by the β-strands of each OB domain and bordered by their connecting loops. Key DNA contacts are made by aromatic or hydrophobic residues that stack or make van der Waals interactions with the ssDNA bases, and basic residues contacting the phosphodiester backbone (**Figure 2b**). Additionally, several polar and acidic residues contact DNA bases. Strikingly, Rpa3 does not contact ssDNA in the structure of PabRPA. The DNA-binding properties of each subunit of PabRPA were tested individually by using SPR. While both Rpa1 and Rpa2 individual subunits are capable of binding ssDNA, no interaction was observed for the Rpa3 subunit (**Supplementary figure S4c**). This property is shared with the eukaryotic RPA where Rpa3 plays a structural role at the heart of the trimerization core but is not primarily involved in ssDNA binding (Bochkareva et al., 1998; Brill and Bastin-Shanower, 1998; Sibenaller et al., 1998).

Comparing the structures of ssDNA-bound PabRPA and the eukaryotic RPA from a fungus *Ustilago maydis* (Fan and Pavletich, 2012) also reveal intriguing differences (**Supplementary figure S5**). Indeed, although the chemical nature of the interactions and the overall ssDNA pathway through RPA are conserved, the DNA-binding grooves of their OBs are substantially different in Archaea and Eukarya. In the structure of PabRPA, basic residues of the β3-β4 helical insertion in OB-2 contact the DNA phosphodiester backbone and delineate a crevice that is narrower than in the structure of the eukaryotic RPA (**Figure 2b & Supplementary figure S5a**). The helical insertion also exists in eukaryotes but adopts a different conformation and is devoid of conserved basic residues (**Supplementary figure S5a**), thereby suggesting that it may be an evolutionary relic derived from an archaeal ancestor. DNA contacts are also substantially different at the level of OB-3 (OB-D in Eukarya). In PabRPA, an archaeal-specific β-hairpin is inserted within the α1-β1 connecting loop (266-311) and makes extensive contacts with ssDNA (**Figure 2b & Supplementary figure S2**). This is particularly interesting as we show later that this loop acts as a molecular switch, which promotes a tetrameric super-complex of PabRPA in absence of ssDNA. On the other hand, the contacts between the OB-2 Zn-finger domain and ssDNA are remarkably conserved between Archaea and Eukarya. Indeed, in the vicinity of their Zn-fingers domains, the interacting residues and the two neighboring bases are perfectly superimposable.

### Cryo-EM structure of the full-length PabRPA bound to short ssDNA

We collected a cryo-EM SPA (single-particle analysis) dataset of a purified complex of the full-length PabRPA and a 35 nucleotide-long poly-dT ssDNA (dT35) (**Figure 2c and Supplementary figure S6**). Given the small size of this protein-nucleic acid complex (96 kDa) and its intrinsic flexibility due to the modular architecture of RPA, our best 3D reconstruction is limited to a global resolution of 6.3 Å (**Supplementary table S2**). However, the 3D reconstruction shows distinguishable structural features, such as the trimeric helical bundle, and individual OB domains that correspond to the trimerization core. Yet, the PabRPA DNA-bound Tri-C can be fitted into the EM density map by using rigid-body refinement (**Supplementary figure S6**). The refined map, as well as the 2D-class averages, show additional density for OB-1 that is positionally flexible. This additional density also shows connectivity to the Tri-C, which can be accounted for by ssDNA traversing the DNA-binding grooves of the OB-2 and OB-1 domains. The single-stranded DNA region across OB-1 and OB-2 was modelled by superimposing PabRPA OB-1 onto the DNA-bound crystal structure of *Homo sapiens* OB-A (Bochkarev et al., 1997) (**Figure 2c & Supplementary figure S5b**). Comparisons with our PabRPA apo crystal structure reveal that OB-1 and OB-2 are positioned differently in the DNA-bound cryo-EM reconstruction, adopting a more extended conformation than in the crystal structure. Electrostatic potential surface analysis reveals that repositioning the OB-1 domain does extend the DNA-binding channel from the trimerization core to the cooperativity module, thereby facilitating the passage of ssDNA from one module to the other.

### Structural basis for the assembly of multiple PabRPA molecules on ssDNA

*In cellulo*, RPA binds and protects much longer ssDNA tracts than the ones used in the assays described above. The ability of PabRPA to bind to long ssDNA substrates was initially assessed by using negative-staining electron microscopy (**Figure 3a**). By using a concentration of PabRPA that accounts for the binding of one PabRPA to ∼30 nucleotides, PabRPA formed nucleoprotein filaments that fully coated a 6.4kb pM13 single-stranded plasmid, showing that purified PabRPA is capable of self-association on long ssDNA. Then, a cryo-EM structural analysis of multiple RPA molecules bound to DNA was undertaken, by using a poly-dT100 oligonucleotide to form an RPA/ssDNA complex containing multiple RPA molecules (**Supplementary figure S7 & Supplementary table S2**). Our best reconstruction was determined at a subnanometer overall resolution of 8.2 Å showing two central PabRPA complexes sandwiched by two additional molecules that are only partially defined in the cryo-EM map (**Figure 3b**). Distinguishable structural features, such as the trimeric helical bundle, and individual OB domains can be observed for the two central molecules in the 2D-class averages, as well as in the refined 3D model. Docking the model of the PabRPA/dT35 complex into the PabRPA/dT100 cryo-EM reconstruction shows that adjacent RPA complexes are connected through an interaction between the Rpa1 OB-1 domain of one RPA complex with the trimerization core of one downstream complex (**Figure 3c & Supplementary movie 2)**. However, the exact positioning and orientation of OB-1 with respect to the trimerization core of the neighboring PabRPA complex could not be determined confidently.

**Figure 3:**
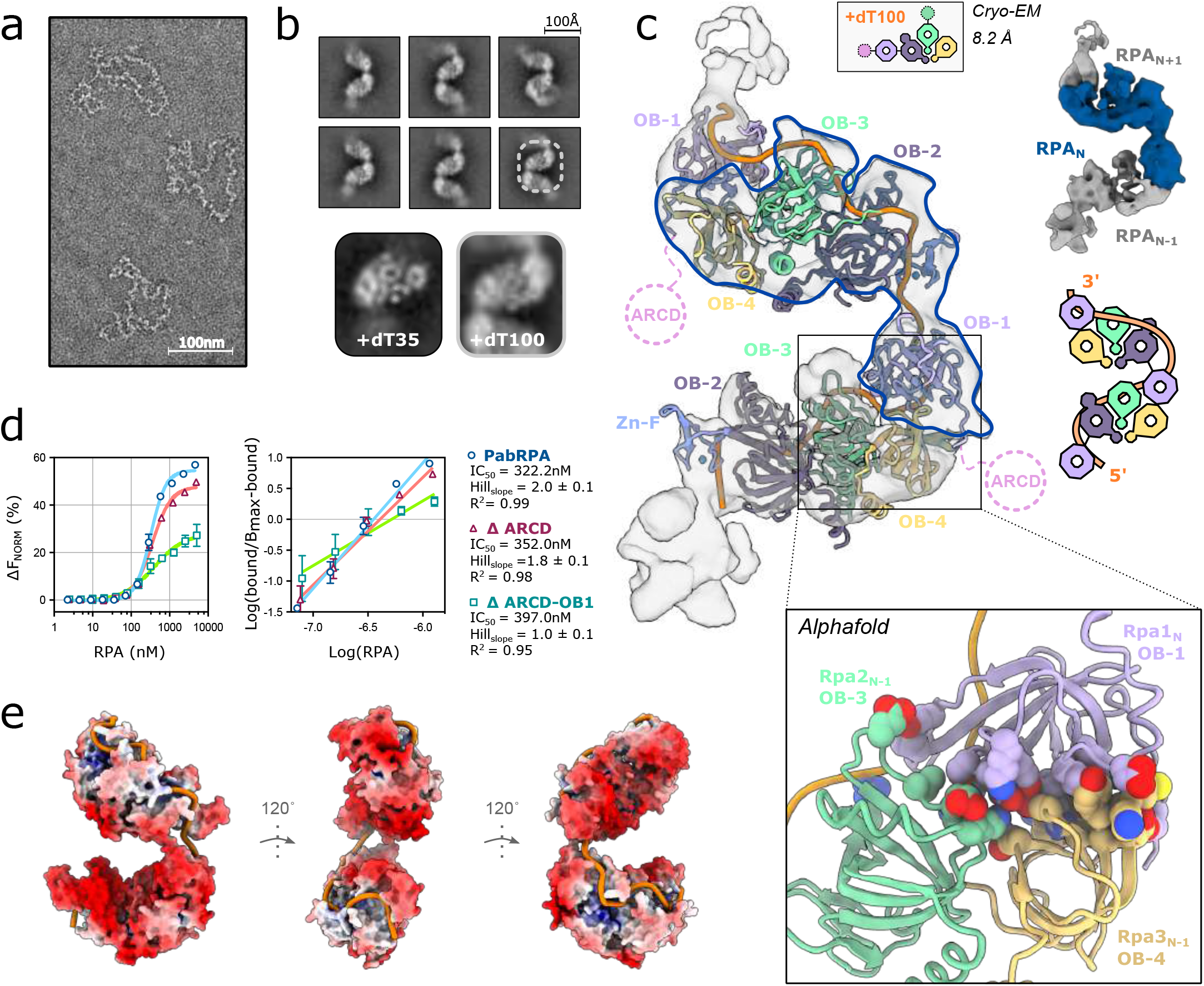
Structural basis for the assembly of multiple PabRPA molecules on ssDNA. **a**. Negative staining microscopy of PabRPA bound to M13 ssDNA plasmid. **b**. Cryo-EM 2D classes of PabRPA bound to a poly-dT100. **c**. Cryo-EM structure of PabRPA bound to a poly-dT100 displayed as backbone traces with a focused view on the Tri-C(n)/OB-1(n+1) AlphaFold model used for model building. The cryo-EM map is contoured at a level of 6rmsd **d**. Biophysical assessment of RPA full-length and cooperativity-module truncated constructs binding onto long ssDNA. Microscale thermophoresis binding curves of RPA heterotrimeric complexes interacting with ssDNA, using wild-type PabRPA (blue), ΔARCD mutant (red) or ΔARCD-OB-1 mutant and a Cy5-labeled poly-dT100 ssDNA substrate. EC_50_ values were calculated using the Hill-equation. Hill coefficients were calculated from the Hill-plot transform of the binding data. **e**. Electrostatic potential surface analysis showing that PabRPA forms a continuous positive groove along the DNA path (orange).

To explore the molecular details of this interaction, we ran an AlphaFold prediction of the protein/protein interactions between two individual PabRPA complexes. The prediction was obtained with a very high level of confidence, predicting that OB-1 contacts simultaneously OB-3 and OB-4 from the adjacent trimerization core (**Figure 3c & Supplementary figure S8a**). AlphaFold predicted protein complex can be nicely fitted in the cryo-EM volumes. DNA was modelled according to our ssDNA-bound crystal and cryo-EM structures of PabRPA (**Figure 2**). Strikingly, the interaction between OB-1(n) and OB-3(n-1) brings together the 3’-end and 5’-end of the modelled ssDNA from two neighboring RPA complexes. The surface electrostatic potential reveals a continuous positively charged cleft that can guide ssDNA from one RPA molecule to the other. Thus, PabRPA nucleoprotein filaments form a polar and continuous groove that efficiently binds and protect ssDNA from nucleases (**Figure 3e & supplementary movie 2**).

The role of the PabRPA cooperativity module was further investigated by using microscale-thermophoresis (MST), an approach that was successfully used to study DNA-binding cooperativity of the yeast RPA (**Figure 3d**) (Yates et al., 2018). As expected, plotting as a linear Hill plot the ssDNA-binding data of full-length PabRPA on long ssDNA (dT100) shows a Hill coefficient of 2.0, thereby confirming that it cooperatively binds to ssDNA. However, plotting ssDNA-binding data of a PabRPA construct deleted from its entire cooperativity module shows a Hill coefficient of 1.0, indicating that this truncated PabRPA construct no longer binds ssDNA cooperatively. Thereby, the critical role of this module in DNA binding cooperativity has been established both by the Cryo-EM structure and by MST. The role of the Archaea-specific domain ARCD was also investigated by using MST. Interestingly, deleting ARCD impaired only partially the binding cooperativity of PabRPA (Hill coefficient of 1.8), indicating that OB-1 plays a more prominent role in cooperativity than ARCD does **(Figure 3d)**. Furthermore, ARCD was not visible in the density of the PabRPA/dT100 cryo-EM reconstruction. This suggests that it participates in PabRPA activity by pre-connecting PabRPA complexes together before they bind DNA but becomes dispensable once the complex is bound to DNA (**Supplementary figure S8b**). Consistently, we verified that deleting ARCD does not alter the DNA-binding affinity of PabRPA to a dT35 short ssDNA suggesting that ARCD is involved in protein/protein interactions rather than protein/DNA interactions (**Supplementary figure S8c**).

### PabRPA forms a tetrameric supercomplex that dissociates upon binding to ssDNA

Unexpectedly, aside from assessing the coating of ssDNA by PabRPA, we also discovered that PabRPA specifically assembles as a tetrameric supercomplex in the absence of DNA by using SEC-SLS (**Figure 4a**). SAXS measurement confirmed that PabRPA oligomerizes spontaneously in solution, forming a compact structure with a radius of gyration (Rg) of 57 Å and a Dmax of 207 Å (**Supplementary figure S9a**). Upon addition of a poly-dT25 ssDNA substrate, the PabRPA tetrameric form dissociates into a smaller complex. To decipher the molecular basis of PabRPA oligomerization in the absence of DNA, the structure of the PabRPA tetrameric supercomplex was determined by cryo-EM at 3.35 Å resolution (**Figure 4b & Supplementary figure S10**). The PabRPA tetrameric supercomplex forms from two dimers (AB and CD) arranged into a C2-symmetrical assembly, which interact through contacts involving their trimerization cores. Interactions within the AB and CD dimers involve their OB-3 domains, while the two dimers are held together by interactions involving their OB-4 domains (**Figure 4c**). PabRPA monomers interact specifically to form the tetramer, burying a great deal of exposed protein surface (2076 Å^2^ per monomer). These observations suggest that assembly of the PabRPA tetrameric complex is thermodynamically favorable in the absence of ssDNA. To evaluate the influence of temperature, the light scattering experiment was repeated at 65°C to get closer to the natural growth temperature of *P. abyssi*. At this temperature, the tetrameric state of PaBRPA was maintained and the intrinsic viscosity increased from 5.0ml/g at 20°C to 5.7ml/g at 65°C showing no change in the overall compact structure of the tetramer but a small gain in flexibility (**Supplementary figure S9b**).

**Figure 4:**
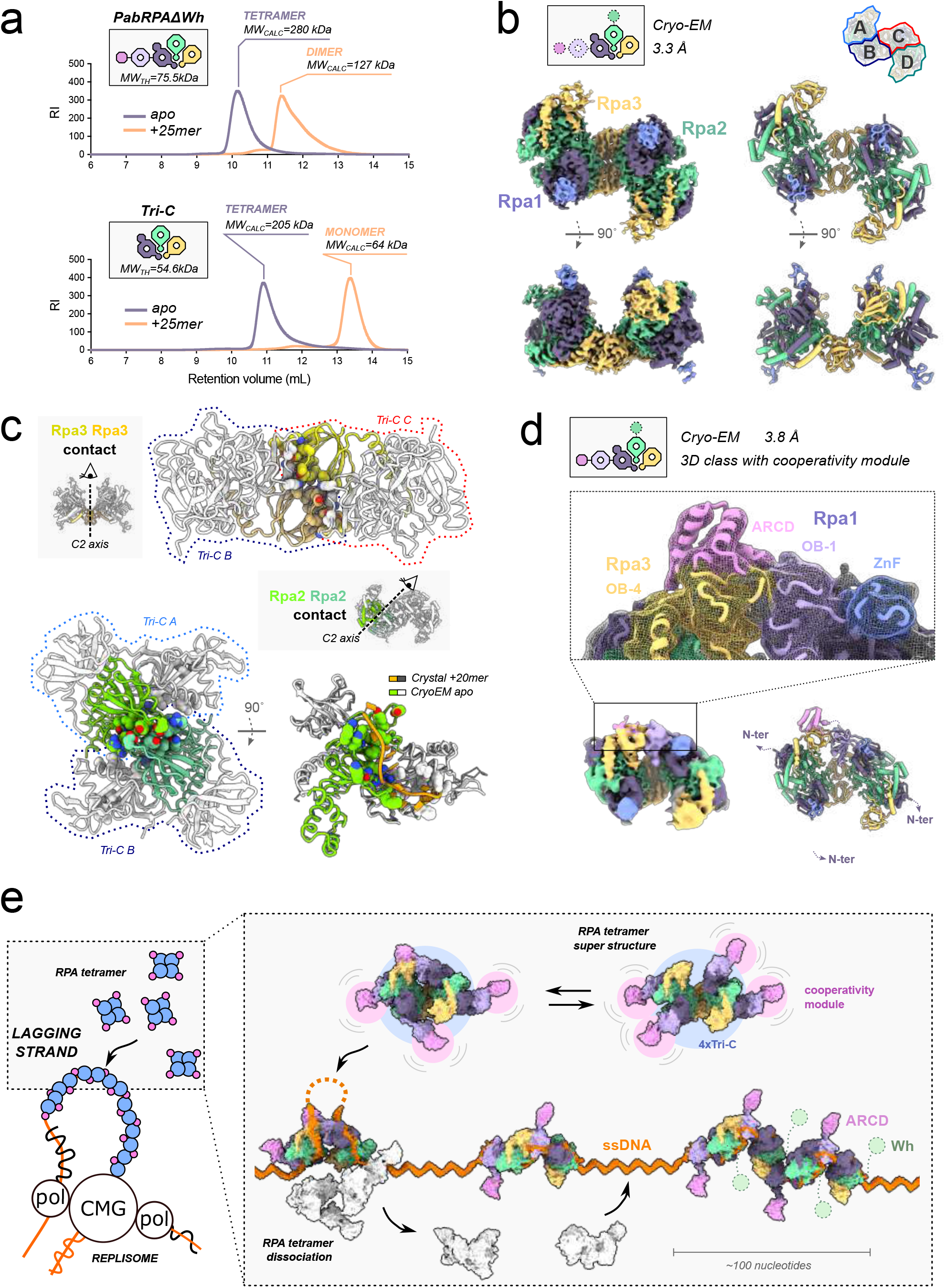
PabRPA assembles as a tetrameric super-complex that dissociates upon binding to ssDNA. **a**. SEC-SLS characterization of PabRPA and PabRPA Tri-C in presence or absence of poly-d25T ssDNA. The theoretical (MW_th_) and calculated molecular weights (MW_calc_) for each complex are given in kDa. **b**. 3.35 Å cryo-EM structure of the tetrameric PabRPA super-structure. **c**. Focused view on critical contacts within the PabRPA tetrameric assembly. **d**. 3D class showing contacts between the cooperativity module and Rpa3 of two neighboring PabRPA molecules within the tetrameric super-structure. **e**. Proposed mechanism for the self-association of PabRPA on ssDNA.

This oligomerization mechanism is strikingly different from the one observed in the DNA-bound PabRPA filaments, which form upon contacts between the cooperativity module and the trimerization core of adjacent RPA molecules. Yet, oligomerization of PabRPA molecules in the absence of ssDNA seems to be independent of the cooperativity module, which was not visible in the cryo-EM reconstruction and appeared to be flexible. SEC-SLS experiments were reproduced in presence or absence of ssDNA by using a truncated version of PabRPA that encompasses only its trimerization core, and lacks the cooperativity module (**Figure 4a**). Consistently, the trimerization core was shown to form a tetramer in solution, and a clear shift into a smaller complex was induced by adding dT25 ssDNA, thereby demonstrating that assembly/dissociation of the PabRPA tetrameric supercomplex is modulated through binding of ssDNA to the Tri-C. It is striking that the OB-3 DNA binding groove is completely occluded in the PabRPA tetrameric supercomplex (**Figure 4c & Supplementary movie S3**). Indeed, many interfacial residues that stabilize the PabRPA tetrameric assembly are also found to bind ssDNA, rationalizing why addition of DNA competes with these intermolecular interactions and promotes dissociation of the PabRPA tetrameric supercomplex. Interestingly, most contacts within the AB and CD dimers involve the α1-β1 β-hairpin of OB-3, which also make extensive contacts with ssDNA (**Figure 2b & Supplementary figure S5**). Thus, the β-hairpin plays a dual role: it drives oligomerization of PabRPA in the absence of DNA, but contacts extensively the substrate in the presence of DNA. Therefore, we hypothesize that this archaeal-specific β-hairpin acts as a molecular switch that promotes a tetrameric super-complex of PabRPA in the absence of ssDNA, contributes to binding the DNA substrate and dissociate the tetramer in the presence of DNA.

Finally, by using a subset of the dataset (**Supplementary figure S10**), we determined an asymetric 3.8 Å cryo-EM structure of the PabRPA tetrameric supercomplex showing the interaction between the cooperativity module of one RPA complex and the OB-4 domain of another RPA (**Figure 4d**). Remarkably, the ARCD/OB-4 interface observed in this new cryo-EM structure matches the one observed in the PabRPA apo crystal structure (**Figure 1b**). This new structure shows how the archaeal-specific ARCD connects two RPA molecules in a conformation that is compatible with their adjacent assembly onto ssDNA.

## DISCUSSION

In all forms of life, coating and protecting exposed ssDNA from nucleases is essential (Oliveira, 2021). In Eukarya, RPA has been shown to be an abundant multi-domain heterotrimeric protein complex, which is essential to all DNA processing events in DNA replication, recombination and repair (Bain et al., 2018; Brosey et al., 2013; Dueva and Iliakis, 2020). Here, we explored the quaternary structure of PabRPA and the conformational changes that occur upon DNA-binding.

### Self-association ssDNA-binding mechanism of PabRPA

We have extensively characterized the RPA from *P. abyssi*, a hyperthermophilic archaeon, and determined three crystal structures and four cryo-EM structures of PabRPA in different states. By using an integrative approach, we uncovered an unanticipated mechanism of self-association of PabRPA on ssDNA (**Figure 4e**). These structures reveal for the first time how ssDNA is handed over from one RPA molecule to the other. In its apo form, PabRPA assembles into tetrameric surpercomplexes of four RPA protomers, whose association is mediated by interactions within OB domains of their trimerization cores (**Figure 4e**). Additional interactions are observed within the cooperativity module and the trimerization cores of two neighboring RPA molecules. Furthermore, the Archaea-specific ARCD domain of one PabRPA molecule has been shown to bind specifically to the OB-4 domain of an adjacent molecule, in the crystal and cryo-EM structures of PabRPA, which were determined in their apo forms. This interaction allows pre-assembly of PabRPA molecules in a conformation that is pre-determined to efficiently coat ssDNA, once the tetramer is dissociated. PabRPA primarily binds ssDNA through its trimerization core, which triggers dissociation of the tetrameric supercomplex. Binding to ssDNA also induces a conformational change within the cooperativity module, untightening the interaction between ARCD(n+1) and OB-4(n) and promoting the binding of the OB-1(n+1) domain to the OB-3(n) and OB-4(n) domains of the neighboring PabRPA complex. This interaction is essential for handing over ssDNA between two adjacent RPA protomers, resulting in coating and protection of ssDNA regions. This cooperative binding mechanism enables the rapid and efficient coating of ∼100 nucleotides per tetramer of PabRPA. The Rpa2 WH domain does not contribute to stabilization of the PabRPA/ssDNA filaments, but instead is ideally positioned to recruit other protein factors involved in DNA metabolism (**Figure 4e**).

### Comparison with eukaryotic RPA

The structures of PabRPA show three essential features conserved in eukaryotes: a trimeric core, a module that promotes cooperative binding to ssDNA and a flexible WH domain, as well as Archaea-specific additional domains. The proposed DNA-binding mechanism of PabRPA displays profound similarities with the dynamic model proposed for the yeast RPA (Yates et al., 2018). Indeed, both studies on PabRPA and yeast RPA support a linear arrangement of ssDNA within RPA (Yates et al., 2018), which differs with the compact horse-shoe conformation that was observed in the crystal structure from *U. maydis* (Fan and Pavletich, 2012). PabRPA possesses a trimerization core and a cooperativity module, which show distinct DNA-binding properties. While the trimerization core associates with ssDNA in a high-affinity stable complex, the cooperativity module binds ssDNA with much lower affinity and higher dissociation kinetics. These findings echo earlier structural, biochemical and biophysical studies showing that the eukaryotic RPA can associate with ssDNA in different modalities, a low-affinity mode involving the Rpa1 N-terminal OB domains and a high-affinity compact mode involving all four major OBs (Arunkumar et al., 2003; Brosey et al., 2015, 2013)(Blackwell and Borowiec, 1994; Fan and Pavletich, 2012). Moreover, multiple RPA molecules assemble on ssDNA by using similar protein-protein interactions in *P. abyssi* and *S. cerevisiae*. Yet, the OB-4(n)/OB-1(n+1) protein-protein interaction in PabRPA (OB-E(n)/OB-A(n+1) in eukaryotes, respectively) plays a critical role when multiple RPAs are associated with ssDNA. In PabRPA, OB-1 (OB-A in eukaryotes) has been shown to be part of a cooperativity module that interacts with the trimerization core of a downstream RPA, thereby enabling the passage of ssDNA across adjacent RPA molecules. Similarly, structural and biophysical studies support that a similar OB-A/OB-E interaction also exists in the yeast RPA and is required for promoting RPA-RPA complexes (Yates et al., 2018). In addition, the OB-4 domain of PabRpa3 does not bind DNA but is a dedicated platform for protein-protein interactions, a property that is shared with the eukaryotic RPA (Bochkareva et al., 1998; Brill and Bastin-Shanower, 1998; Sibenaller et al., 1998). Similarly to previous study on eukaryotic RPA(Yates et al., 2018), the PAbRPA2-WH domain remains unseen in both crystallography and cryo-EM structures, most likely due to its highly flexible attachment to the RPA trimeric core. This conserved flexibility is probably a key element that guaranties its accessibility.

A recent structure of the RPA-like human CST complex revealed a decameric assembly bound to telomeric DNA (Lim et al., 2020). In Archaea, the PabRPA tetrameric supercomplex may play a regulatory role by clustering multiple PabRPA complexes, which can rapidly and efficiently bind to exposed ssDNA tracts. Our work strongly suggests that association/dissociation of the PabRPA tetrameric supercomplex is controlled by a β-hairpin molecular switch, which is located in the N-terminal region of Rpa2. However, multimerization of RPA, such as the PabRPA tetrameric supercomplex, has not been reported in eukaryotes. Yet, the equivalent N-terminal region of Rpa32 in eukaryotes hosts important regulatory phosphorylation sites that control the DNA damage signaling and repair activities of RPA (Dutta and Stillman, 1992; Kõivomägi et al., 2011, p. 1; Maréchal and Zou, 2015; Pan et al., 1994; Stephan et al., 2009). This observation highlights the key regulatory roles that the N-terminal region of Rpa2 (Rpa 32 in eukaryotes) play in DNA-binding activities of RPA. PabRPA also shows intriguing specificities, such as the Archaea-specific ARCD domain, which allows pre-assembly of RPA molecules in a conformation that is pre-determined to efficiently coat ssDNA. Strikingly, while this domain is widely distributed among Archaea, it is absent in eukaryotes. Instead of ARCD, the eukaryotic Rpa1 N-terminal extension includes an extra OB domain that is dedicated to protein-protein interactions, and is linked to OB-A by a 50 amino-acid long linker with no predicted secondary structures. Aside from its role in connecting RPA protomers together, one may hypothesize that ARCD also recruits genome maintenance factors once PabRPA is bound to DNA.

While the newly characterized ARCD domain is widely distributed across Archaea, being present in nearly all Rpa1 and Rpa1-like subunits, the Rpa2 C-terminal helicoidal extension and the β-hairpin molecular switch of PabRPA may be specific to particular groups of Archaea. Interestingly, Archaea possess a highly differentiated distribution of RPA (MacNeill, 2021; Taib et al., 2021). *Pyrococcus furiosis* and *Thermococcus kodakarensis* encode three RPA subunits, which form a heterotrimer with high affinity for ssDNA (Komori and Ishino, 2001; Nagata et al., 2019). *Methanosarcina acetivorans* also encode three RPA-like SSBs, but they do not interact, and all form homodimers (Robbins et al., 2004). Non-canonical RPA molecules, which display intriguing DNA-binding properties, have also been characterized in *Haloferax volcanii* but the exact stoichiometry of their complexes is unknown (Stroud et al., 2012). In Archaea, RPA displays a wide range of domain architectures with varying numbers of OB domains. For example, one or three additional OB domains are respectively inserted between the OB-1 and OB-2 domains in *M. acetivorans* and *M. jannaschii* Rpa1 subunits (MacNeill, 2021), and this may affect the DNA-binding properties of these RPAs. Therefore, extending this study to RPA from other archaeal species would be of particular interest.

### Evolution of RPA in Archaea and Eukarya

This work constitutes a significant step towards the molecular understanding of the structure and DNA-binding mechanism of RPA, with important implications for the evolution of this vital replication factor in Archaea and Eukarya. In Bacteria, the archetypal SSB is the major single-stranded DNA binding protein (Theobald et al., 2003). It encompasses a single OB domain and assembles into homotetrameric complexes (Meyer and Laine, 1990). Eukaryotes also encode single OB-fold SSBs, but their function is restricted to DNA damage repair, whereas the major ssDNA-binding component of the replisome consists of a heterotrimeric RPA complex (Brosey et al., 2013). In addition to RPA, several RPA-like complexes have evolved to perform various specialized roles, as in the case of the CST (Cdc13-Stn1-Ten1) complex, which is essential for telomer maintenance (Lim et al., 2020; Miyake et al., 2009; Surovtseva et al., 2009). By contrast, Archaea display a patchier distribution of SSB and RPA proteins, with some archaeal lineages encoding either one or both of the two systems (Raymann et al., 2014; Wadsworth, 2001). The genomic neighborhoods of the archaeal RPA genes are also highly variable (**Supplementary figure S3**). Such SSB/RPA distribution patterns prompt questions regarding the emergence of the trimeric RPAs and the provenance of the different eukaryotic ssDNA-binding proteins. These questions are challenging to address using sequence-based comparisons due to small size and high divergence of the OB domains. Instead, we performed here an all-against-all structural comparison of the OB domains from representative SSB and RPA originating from Bacteria, Eukarya and Archaea as well as eukaryotic RPA-like CST complex (**Figure 5a**). The available structures were supplemented with AlphaFold models (Jumper et al., 2021) of additional crenarchaeal and thaumarchaeal SSBs and trimeric RPAs from Asgard archaea, the postulated ancestors of eukaryotes (Liu et al., 2021; Spang et al., 2015). The structures formed three major branches in the structural distance-based tree. The first branch included all bacterial SSBs. In the second clade, thaumarchaeal, crenarchaeal and eukaryotic SSBs grouped with the archaeal and eukaryotic OB-1 domain of Rpa1 (OB-A in eukaryotes), with OB-2 of Rpa1 (OB-C in eukaryotes) forming a sister group to this assemblage. Finally, the third clade included two sister clades corresponding to OB-3 and OB-4 domains of archaeal and eukaryotic Rpa2 and Rpa3, respectively. It was previously noted that it is difficult to distinguish single-OB domain SSBs and Rpa3 (Taib et al., 2021). Structural comparisons clearly distinguish the two sets of proteins and strongly suggest an evolutionary relationship between the archaeal SSBs and the N-terminal OB domain of archaeal and eukaryotic Rpa1 subunits, rather than with Rpa3. These observations are consistent with the evolution of heterotrimeric RPA from a single-OB SSB ancestor, likely within the archaeal branch (Gamsjaeger et al., 2015; Kerr, 2003; Morten et al., 2015). Given the shared non-structured C-terminal extension in bacterial and archaeal SSBs and the ability to oligomerize, it is likely that they have evolved from a common ancestor, which might have functioned already in the last universal cellular ancestor (LUCA). Within archaea, this SSB gave rise to the multidomain Rpa1 through tandem duplication followed by insertion of a Zn-finger subdomain into OB-2 (**Figure 5b**). By contrast, Rpa2 and Rpa3 appear to be structural paralogs, with Rpa2 accruing a characteristic C-terminal WH domain. Interestingly, the PabRPA OB-3 and OB-4 domains respectively share the highest structural similarity with the OB-domains from the Ctc1 and Ten1 subunits of the human telomeric maintenance CST complex. More generally, the OB domains from eukaryotic RPA and RPA-like complexes, such as the CST, grouped with the corresponding domains of archaeal PabRPA-like homologs, including those from Asgard archaea, suggesting that the eukaryotic RPA and its derivatives have been inherited from the archaeal ancestors.

**Figure 5:**
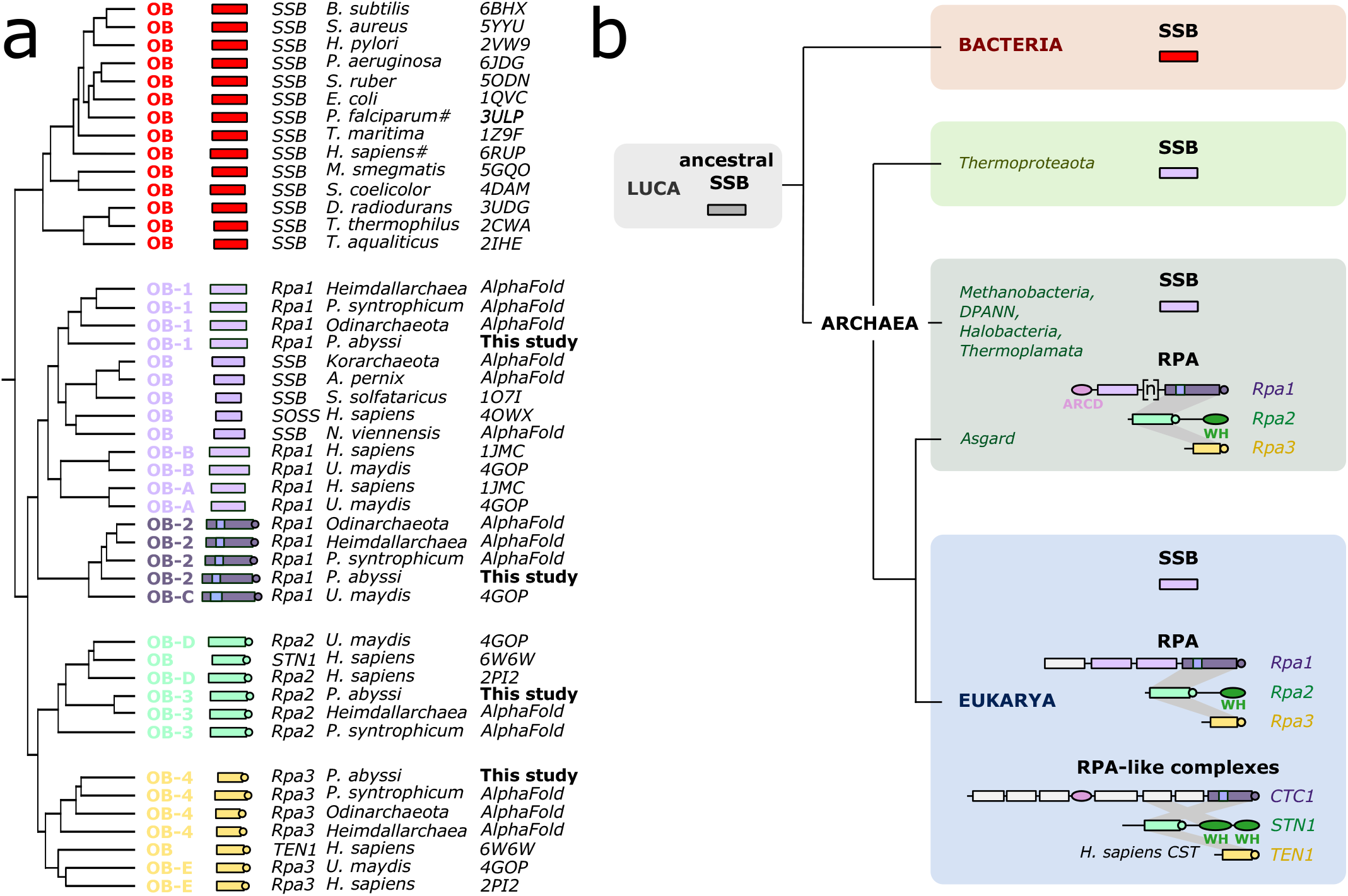
Evolutionary implications for the RPA evolution in Archaea and Eukarya. **a**. Structure-based dendrogram produced by DALI, based on average linkage clustering of the Z-scores of the structural similarity matrix. **b**. Current proposal for the evolutionary history of SSB and RPA in the three domains of life.

RPA, which evolved from its SSB ancestor by accretion of complexity, has acquired the ability to accomplish a broad range of biological functions in DNA replication, recombination and repair (Oliveira, 2021). In addition, this increased level of complexity probably also enabled more sophisticated mechanisms of modulation and regulation of RPA. Indeed, RPA is a protein platform that recruits many different key actors involved in genome maintenance (Brosey et al., 2013). The strong structural similarity observed between PabRPA and the human telomeric maintenance CST complex is also intriguing. Indeed, telomer maintenance in linear genomes are defining features of eukaryotes compared to prokaryotes, which possess circular genomes. Therefore, the inherited ancestral RPA may have provided the primitive eukaryotes with key genome maintenance activities that were required for the emergence of the eukaryotic domain of life.

## MATERIEL & METHODS

### Cloning, protein expression, and purification

The open reading frames (ORFs) of the Rpa1, Rpa2 and Rpa3 genes from *P. abyssi* were optimized and synthesized by GeneArt (Thermo Fisher). For individual subunit expressions, ORFs were inserted into the pRSFduet(+) multiple cloning site 1 with a TEV-cleavable N-terminal 14-His tag. For PabRPA complexes co-expressions, ORFs were inserted into the pRSFduet(+) multiple cloning site 1 as a polycistronic Rpa3-Rpa2-Rpa1 construct separated by ribosome binding sites (RBS), with a TEV-cleavable N-terminal His14-tagged Rpa3 fusion protein. The folowing RPA isoforms were cloned from the pRSFduet(+) constructs using the Q5 site directed mutagenesis kit (New England Biolabs): PabRpa1 cooperativity module (Rpa1_1-193_), PabRPA trimerization core Tri-C (Rpa1_193-358_/Rpa2_1-179_/Rpa3), PabRPA ARCD (Rpa1_64-358_/Rpa2/Rpa3), PabRPA ΔARCD-OB-1 (Rpa1_193-358_/Rpa2/Rpa3), PabRPA H (Rpa1/Rpa2_1-179_/Rpa3).

Proteins were expressed in BL21 Star (DE3) strain from *E. coli* (Invitrogen) at 37°C in LB medium supplemented with 100 μg/mL of kanamycin. Recombinant protein expression was induced by adding 0.25 mM IPTG. Cells were then incubated overnight at 20°C, collected by centrifugation, resuspended in buffer A (0.02 M Na-HEPES at pH 8, 0.5 M NaCl, 0.02 M imidazole) supplemented with complete EDTA-free protease inhibitors (Roche), and lysed with a Cell-Disruptor. Lysates were then heated for 10 min at 60°C and centrifuged 30 min at 20000 g. PabRPA purifications were performed using a three-step protocol including nickel affinity, anion exchange and size exclusion chromatography. The clear cell lysate was loaded onto 5 mL HisTrap columns (Cytiva) connected to an ÄKTA purifier (Cytiva). Elutions were performed using a linear gradient of imidazole (buffer B, 0.02 M Na-HEPES at pH 8, 0.5 M NaCl, 0.5 M imidazole). Protein fractions were combined, dialyzed in buffer C (0.02 M Na-HEPES pH 8, 0.1 M NaCl), loaded onto 5 ml HiTrap Q FF columns (Cytiva) and eluted with a linear gradient, by mixing buffer C with buffer D (0.02 M Na-HEPES pH 8, 2 M NaCl). Depending on the applications, the 14-His tag was removed following an overnight TEV-protease cleavage. Purifications were finally polished using exclusion-size chromatography in buffer E (0.02 M Na-Hepes pH 8, 0,15 M NaCl) on a Superdex 200 10/300 (Cytiva).

### Crystallization

Crystallization trials were performed using the hanging drop vapor diffusion technique in 2 μL drops (1:1 reservoir to protein ratio) equilibrated against 500 μL of reservoir solution. PabRPA crystals were obtained at 18°C in 0.1 M Tris pH 8.5, 1.2 M ammonium sulfate with a protein solution at 10 mg/mL. After 72 hours, crystals were transferred in a dehydration solution containing the original mother liquor supplemented with 25% glycerol and incubated for 1 hour at 4 °C before flash-freezing in liquid nitrogen. PabRpa1 cooperativity module crystals were obtained at 18°C using a crystallization solution made of 0.01 M NiCl2, 20% w/v PEG MME 2K and 0.1 M Tris pH 8.5 with a protein solution at 40 mg/mL. Crystals were then directly flash-frozen in liquid nitrogen without additional cryoprotection. The PabRPA Tri-C/dT20 complex was reconstituted by mixing the protein solution at 0.5 mg/mL with a 1.2X excess of poly-dT20 ssDNA (Eurogentec) in buffer E. The mixture was then concentrated ∼30 times to obtain a final OD_280nm_ value of 40. Crystals were grown at 4°C in 10% w/v PEG 8K, 0.1 M imidazole pH 8 and 0.2 M calcium acetate, and cryoprotected with 25% ethylene glycol. X-ray data were collected at the SOLEIL synchrotron on beamlines PX1 and PX2.

### X-ray data collection, processing, model building and refinement

Diffraction data collection and refinement statistics are given in **Supplementary table S1**. Crystallographic data were collected on the PROXIMA-1 (Chavas et al., 2021) beamlines at Synchrotron SOLEIL (Saint-Aubin, France) and processed with XDS (Kabsch, 2010) through XDSME (https://github.com/legrandp/xdsme) (Legrand, 2017). The strong diffraction anisotropy was corrected using the STARANISO program (Tickle et al., 2018). Initial phases for the apo-PabRPA crystal structure were obtained by Zn-SAD. The diffraction data from 3 crystals collected at the peak absorption wavelength of zinc K-edge (1.2827 Å) were merged to obtain a highly redundant dataset (>60). A unique zinc site was found using the SHELXC/D (Sheldrick, 2008) from which phases were calculated with PHASER (McCoy et al., 2007) and then improved by density modification with the PARROT program (Cowtan, 2010). Thus, a first experimental electron density map at 4.5 Å resolution could be obtained in which an initial model could be assembled and completed using COOT (Emsley et al., 2010) combining the N-terminal region of Rpa1 previously determined and other partial models like the zinc finger. Later, this initial model could be improved with the help of AlphaFold (Jumper et al., 2021) predictions, and refined using a dataset reaching 3.7 Å in the best diffracting direction. The procedure proposed by Terwilliger (Terwilliger et al., 2022) was exploited to iteratively improved AlphaFold models using experimental information. Crystal structures of the PabRpa1 cooperativity module and the PabRPA Tri-C/d20T complex were determined by molecular replacement with MOLREP (Vagin and Teplyakov, 1997) using the previously determined apo structure of PabRPA. All refinements were conducted with the BUSTER program (Bricogne, G. et al., 2017) using TLS motion groups and COOT was used for model reconstruction. In the case of PabRPA-apo and PabRPA Tri-C/d20T, local structure similarity restraints (LSSR) were used using hybrid target models constructed from a mix of higher resolution structures from the present work and AlphaFold predicted models (Jumper et al., 2021; Terwilliger et al., 2022). Details for all datasets are summarized in **Supplementary table S1**.

### AlphaFold model predictions

AlphaFold model predictions were calculated either using the Google Colab platform and AlphaFold2_advanced form developed by the ColabFold team (Mirdita et al., 2021) (https://colab.research.google.com/github/sokrypton/ColabFold) or using a local installation of ColabFold obtained from the LocalColabFold Github repository (https://github.com/YoshitakaMo/localcolabfold).

### Cryo-EM sample preparation

The PabRPA/dT35 complex was reconstituted by mixing a poly-dT35 ssDNA (Eurogentec) with a 3-fold excess of PabRPA. The complex was then injected on a superdex 200 10/300 in buffer F (0.02 M HEPES pH 8, 0.1 M NaCl and 0.002 M magnesium acetate). 3 μL of the eluted fraction (OD_280nm_=1.5) were applied to glow-discharged quantifoil R2/2 300 gold mesh grids (Electron Microscopy Sciences). After 20 seconds of pre-incubation, the grids were blotted for 6 seconds (force -2) using a Vitrobot Mark IV (ThermoFischer) at 100% humidity and 25 °C. The PabRPA/d100T complex was reconstituted using the same strategy but using a 5-fold excess of PabRPA. Specimens were prepared similarly with quantifoil R2/2 200 copper mesh grids (Electron Microscopy Sciences). For the PabRPA tetrameric supercomplex, 3 μL of purified PabRPA at 0.5 mg/mL were applied to glow discharged Quantifoil R2/2 300 gold mesh grids. The grids were then blotted for 6 seconds (force 0).

### Cryo-EM data acquisition and image processing

Movie acquisition was carried out at the Nanoimaging Core Facility (Institut Pasteur, Paris). Details for all datasets are summarized in table 2. The PabRPA/dT35 dataset was collected on a 200 kV Glacios electron microscope (Thermo Fisher) equipped with a Falcon 3 detector operating in electron counting mode. The PabRPA/dT100 dataset was collected in the same microscope, operating in integration mode. The PabRPA tetrameric supercomplex dataset was collected on a 300 kV Titan Krios electron microscope (Thermo Fisher) equipped with a K3 detector and a bio-quantum energy filter (Gatan).

Motion correction and CTF-estimation of the acquired movies were carried out in Cryosparc v3.3.2 (Punjani et al., 2017). Image processing pipelines for each map are shown in **Supplementary figures S4, S5 & S7**. Blob picking was used to generate initial picks of PabRPA/dT35, which were then filtered by 2D classification and used to train a Topaz picking model (Bepler et al., 2019). Topaz particle picks were used to generate three initial models, which were refined with heterogeneous refinement jobs, including a volume encompassing only the trimeric core. The highest resolution map was refined using non-uniform refinement yielding a 6.24 Å resolution map, and then locally filtered with a lanczos filter. Initial templates of PabRPA/dT100 were generated with blob picking, and then used for template picking. Multiple initial models were generated, and the one that showed the interaction between two RPA complexes was chosen and refined through heterogeneous and non-uniform refinement, achieving 8.24 Å resolution. Initial 2D templates of the PabRPA tetrameric supercomplexes datasets were generated via blob-picking and used to train a Topaz model. Topaz picks were then used to generate three initial models, which were refined using heterogeneous refinement. 3D classification yielded a map containing the cooperativity module, which was refined using non-uniform refinement and locally filtered with a lanczos filter, achieving 3.8 Å resolution. A high-resolution map that did not include the cooperativity module was refined with non-uniform refinement and C2 symmetry to achieve 3.35 Å resolution. Details for all datasets are summarized in **Supplementary table S2**.

### Building and refinement of cryo-EM models

<u>Procedure for the PabRPA/d35T cryo-EM structure:</u> individual crystal structures of PabRPA Tri-C/d20T (PDBid: 8AAS) and PabRpa1 OB-1 (PDBid: 8AA9) were manually placed in the cryo-EM map and subsequently rigid-body fitted in the density with coot (Emsley et al., 2010; Emsley and Cowtan, 2004). Six different rigid-body groups were used: the trimerization bundle, the four individual OB domains and the Rpa1 zinc-finger domain (**Supplementary figure S6f**). ssDNA modeling was guided by superimposing the PabRRPA Tri-C/d20T (PDBid:8AAS) and the human Rpa1 OB-A/ssDNA (PDBid:1JMC) crystal structures. This initial model was further real-space refined by using Phenix (Adams et al., 2011), with a high weight on ideal geometry and restraints on secondary structures. The final map correlation coefficient is 0.63. <u>Procedure for the PabRPA/d100T cryo-EM structure:</u> two identical models of Tri-C(n)/OB-1(n+1) complex predicted by AlphaFold (Evans et al., 2021) were manually placed and rigid-body fitted in the cryo-EM map using coot (**Supplementary figure S7f**). Modelling of the ssDNA substrate was guided by superimposing the PabRPA/d35T cryo-EM structure (PDBid: 8ACE). Finally, the complete model was real-space refined using Phenix, following the same procedure as for the PabRPA/d35T structure (See above). The final map correlation coefficient is 0.64. <u>Procedure for the PabRPA tetrameric supercomplex cryo-EM structure:</u> Four identical models of Pab-RPA Tri-C complex (PDBid: 8AAS) were manually placed and rigid-body fitted in the cryo-EM map using coot. The initial model was then subjected to global real-space refinement program from Phenix using secondary structure restraints. The refined model was further manually inspected and adjusted in coot. <u>Procedure for the PabRPA tetrameric supercomplex cryo-EM structure with the cooperativity module connecting two PabRPA molecules:</u> The model was built by using the same procedure than for the PabRPA tetrameric complex (see above). In addition, Rpa1 ARCD and OB-1 domains crystal structures (PDBid: 8AA9) were manually placed and rigid-body fitted using coot and Phenix. Refinement details for all datasets are summarized in **Supplementary table S2**.

### Structure analysis

Electrostatic calculations were performed by using APBS (Jurrus et al., 2018). Interaction surfaces were calculated by using the PISA webserver (Krissinel and Henrick, 2007). Surface conservations were calculated using CONSURF (Ashkenazy et al., 2016). Structure comparisons were performed by using DALI (Holm, 2020). All figures were prepared using UCSF Chimera X (Pettersen et al., 2021).

### Negative staining microscopy

The PabRPA/pM13 complex was reconstituted by mixing 1 μL of M13mp18 ssDNA (New England Biolabs) at 250 μg/mL with 1 μL of PabRPA at 30 μM (200X excess) in 98 μL of buffer F. 3 μL of the mixture were deposited on glow-discharged carbon-coated copper grids CF400-CU (Electron Microscopy Sciences) and contrasted 3×1 minute in 2% uranyl acetate. Data collection was performed using a Tecnai biotwin T12 (Thermo Fisher) equipped with a LaB6 filament, operating at 120 keV. Images were recorded using an Eagle camera (Thermo Fisher) at a nominal magnification of 49000, using a 3 μm defocus.

### Electromobility shift assay

2.5 pmol of 3’-FAM ssDNA-24mer (5’-GCCTGCAGGTCGACTCTAGAGGAT-3’) or dsDNA-24mer were mixed with increasing amounts of *Pab*RPA in Tris-HCl 30 mM pH 7.5, NaCl 300 mM, 0.5 g/L bovine serum albumin (BSA), 5% Ficoll (20 μL final volume). Reactions were carried out to reach equilibrium at 60°C for 1 h before migration in 1% agarose gel at 4°C under native conditions (90 V, 30 mA). Images were acquired with Typhoon9400 (GE Healthcare).

### Surface plasmon resonance (SPR)

Surface plasmon resonance data have been acquired with Reichert SR7000DC spectrometer equipped with a self-sampling injection system Reichert SR7100. Sensor chips consisted of a glass slide coated by a gold film, the latter being already functionalized by a mixed self-assembled monolayer (SAM) of 90% dithiol aromatic PEG6-COOH and 10% of dithiol aromatic PEG3-COOH. Neutravidin was immobilized onto SAM, in both flow cells, using classical EDC:NHS coupling chemistry. Appropriate amounts of DNA were immobilized onto surfaces by successive injections at 25 μL/min of 1 μM of 5’-biotin-TEG labelled 32-mer ssDNA (5’-TGCCAAGCTTGCATGCCTGCAGGTCGACTCTA-3’), ssRNA (5’-UGCCAAGCUUGCAUGCCUGCAGGUCGACUCUA-3’) or dsDNA (Eurogentec) diluted in buffer E supplemented with 0.05% tween. Binding assays were performed at 25°C by injecting increasing concentrations of protein solutions for 300 s at 25 μL/min. The raw sensograms were processed by double referencing i.e. subtracting both the signals measured on the reference flow cell and the signals measured for blank injections. Resulting sensorgrams were analyzed by Scrubber2.0 (BiolLogic) following several Monte Carlo fits without mass transport limitation of single equilibrium system for PabRPA, Rpa2, Rpa3 and Rpa1/Rpa2 and two-steps equilibrium system for Rpa1.

### Bio-layer interferometry (BLI)

ssDNA-protein binding assays were performed on an Octet RED384 BLI instrument (ForteBio). 3’-biotin-TEG labelled poly-dT35 ssDNA were captured at a concentration of 5 μg/mL for 100 s on streptavidin biosensors. Binding experiments were monitored at 25°C in buffer E supplemented with 0.2 mg/mL BSA (Sigma). Data were analyzed with Prism 9.4. A sample reference with buffer-only was subtracted from all curves. Affinities were determined by fitting the concentration dependence of the experimental steady-state signals, using the following equation Req = Rmax*(RPA)/(Kd + (RPA)).

### Microscale thermophoresis (MST)

The MST measurements were performed at room temperature on Monolith NT.115 Blue/Red (NanoTemper Technologies, GmbH) instrument using standard capillaries. For all experiments, the LED power and IR laser configuration were set to 20% and default, respectively. The MST data were either analyzed with NanoTemper Analysis v2.3, Prism 9.4 or Matlab 2016b software. In all cases, the data were fitted using the Hill equation without any constrains. Each experiment was performed at least three times in buffer E. 5’-Cy5 fluorophore labelled poly-dT100 ssDNA (Integrated DNA technologies) was diluted to a concentration of 200nM and mixed with an equal volume of a serial dilution series of WT or mutants PabRPA and incubated over night at 20°C before loading into MST capillaries.

### Small-Angle X-Ray Scattering (SAXS)

SAXS data of the PabRPA ΔWH complex (50 μl at 1 mg/ml) in buffer E was collected in batch Mode at beamline SWING (synchrotron SOLEIL). The curves were background-subtracted using FOXTROT and analyzed using the ATSAS 3.0.2 software suite. The normalized Kratky was calculated according to (Durand et al., 2010; Pérez et al., 2001) using the value determined by Guinier analysis. P(r) functions were computed from the scattering curves by an indirect transform method in GNOM (Svergun, 1992).

### Molecular mass measurements by size exclusion chromatography coupled to static light scattering detection (SEC-SLS) and viscometry

The oligomerization states of PabRPA H and PabRPA Tri-C at 20 μM in presence or absence of 24 μM poly-dT25 ssDNA (Eurogentec) were determined by size exclusion chromatography coupled to a triple detection (concentration detector: UV detector, refractometer; static light scattering 7°, 90°; viscometer) on a Omnisec resolve/reveal instrument (Malvern Panalytical). The column and detectors were equilibrated with the filtered and degazed running buffer E prior measurement. All proteins were injected (100 μl) and eluted at 0.4 ml/min on a Superdex 200 increase 10/300 GL column (Cytiva). Detections were performed at 20°C or 65°C. External calibration was done with BSA using an injection of 100 μl at 1.4 mg/ml. The refractive index, static light scattering, and the viscosity measurements were processed to determine the mass average molecular mass and the intrinsic viscosity using the OMNISEC V11.10 software (Malvern Panalytical).

## Data avaibility

The PabRpa1 cooperativity module, PabRPA and PabRPA Tri-C/d20T crystal structures were deposited in the Protein Data Bank under accession codes, 8AA9, 8AAJ and 8AAS respectively. The all-atom cryo-EM structure of PabRPA tetramer, as well as the 3D-class showing the cooperativity module are deposited under accession code (PDBid: 8AAH; EMDB-15300) and (PDBid: 8AAL; EMDB-15301) respectively. Finally, C-alpha backbone, nucleic acid phosphate trace and the zinc metal ions atomic coordinates of the PabRPA/d35T and PabRPA/d100T complexes are deposited under the accession codes (PDBid: 8ACE; EMDB-15347) and (PDBid: 8ACJ; EMDB-15350), respectively.

## Acknowledgments

We thank the Platforms for X-ray crystallography (PF6), biophysics, and nanoimaging in the Institut Pasteur. We acknowledge SOLEIL for provision of synchrotron radiation facilities and the staff of beamline PROXIMA-1 (Saint-Aubin, France). We gratefully acknowledge the financial support of ANR (Grants ARCHAPOL and ARCHAPRIM). C.M. was funded by a postdoctoral Pasteur-Roux-Cantarini fellowship. M.M.C was funded by a postdoctoral FRM fellowship (Fondation pour la Recherche Médicale). We would like to thank Sebastien Brûlé (Molecular Biophysics Platform, C2RT, Institut Pasteur) for helping us with SEC-SLS, and Olympe Vaillant for preparing the RPA topology diagram. We also would like to thank Dr. Christophe Crézé for performing the EMSA experiments.

## SUPPLEMENTARY FIGURE AND TABLE LEGENDS

**Figure S1:**
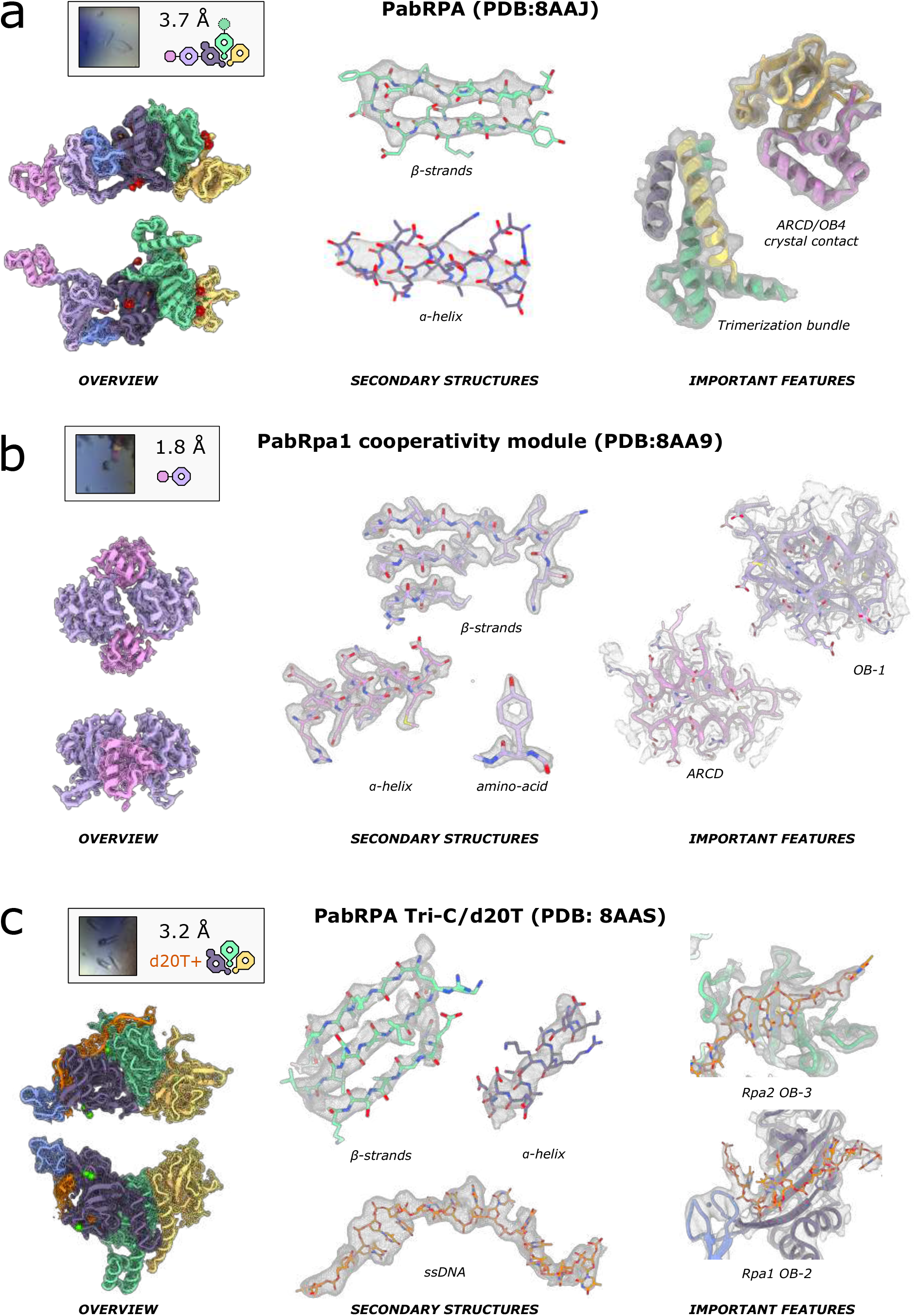
Electron density in representative regions of the PabRPA crystal structures. PabRPA (**a**), PabRPA Rpa1 cooperativity module (**b**) and PabRPA Tri-C/20dT (**c**) crystal structures are represented as Cα traces in the left panels. Right panels show close views of functionally important regions of PabRPA. Main chain and side chains atoms are represented as Cα traces and sticks, respectively. The gray mesh is the maximum likelihood 2mFo-DFc electron density map contoured at a level of 1 σ.

**Figure S2:**
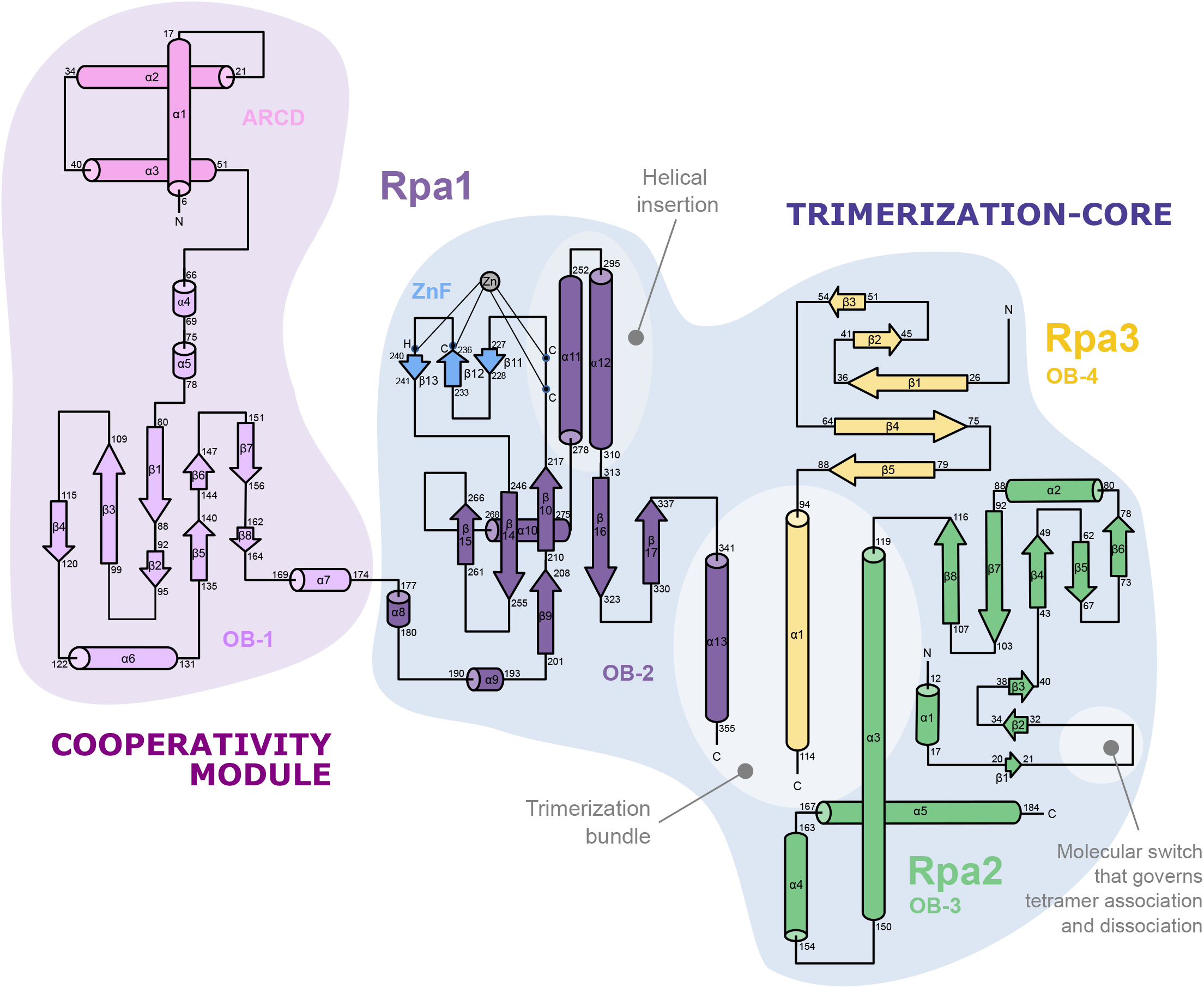
Topological diagram of PabRPA. Two-dimensional diagram of protein topology for three subunits of PabRPA.

**Figure S3:**
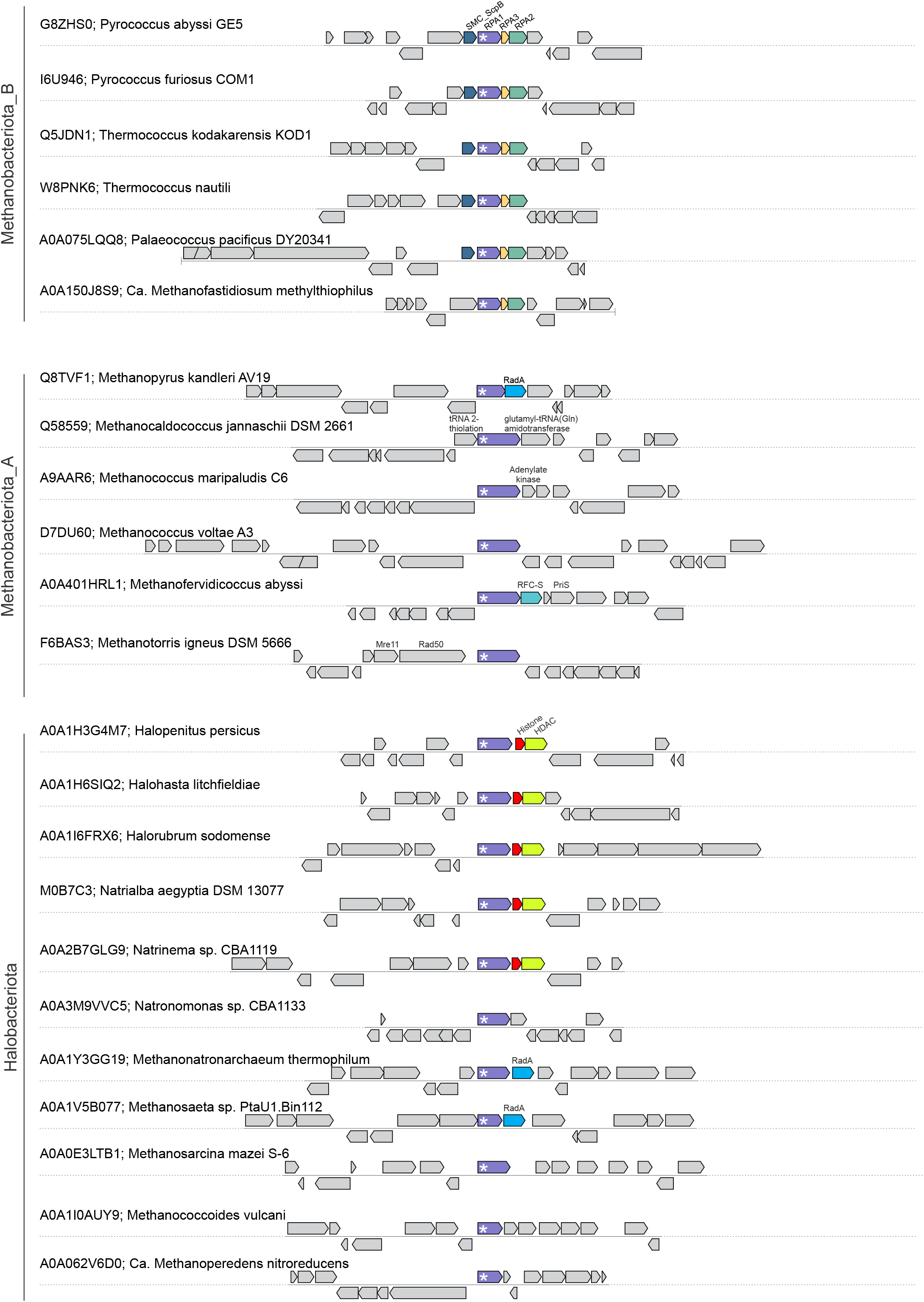
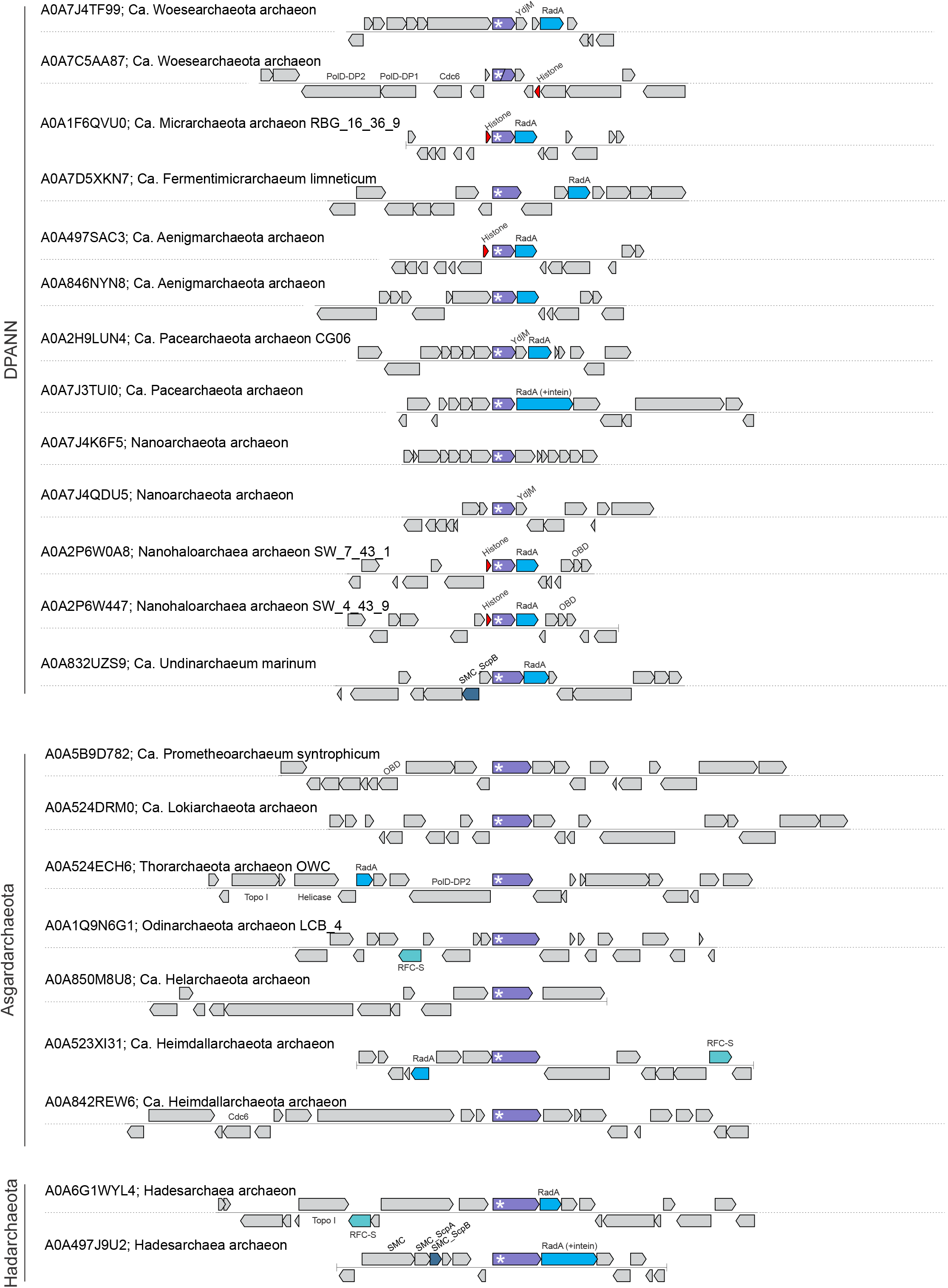
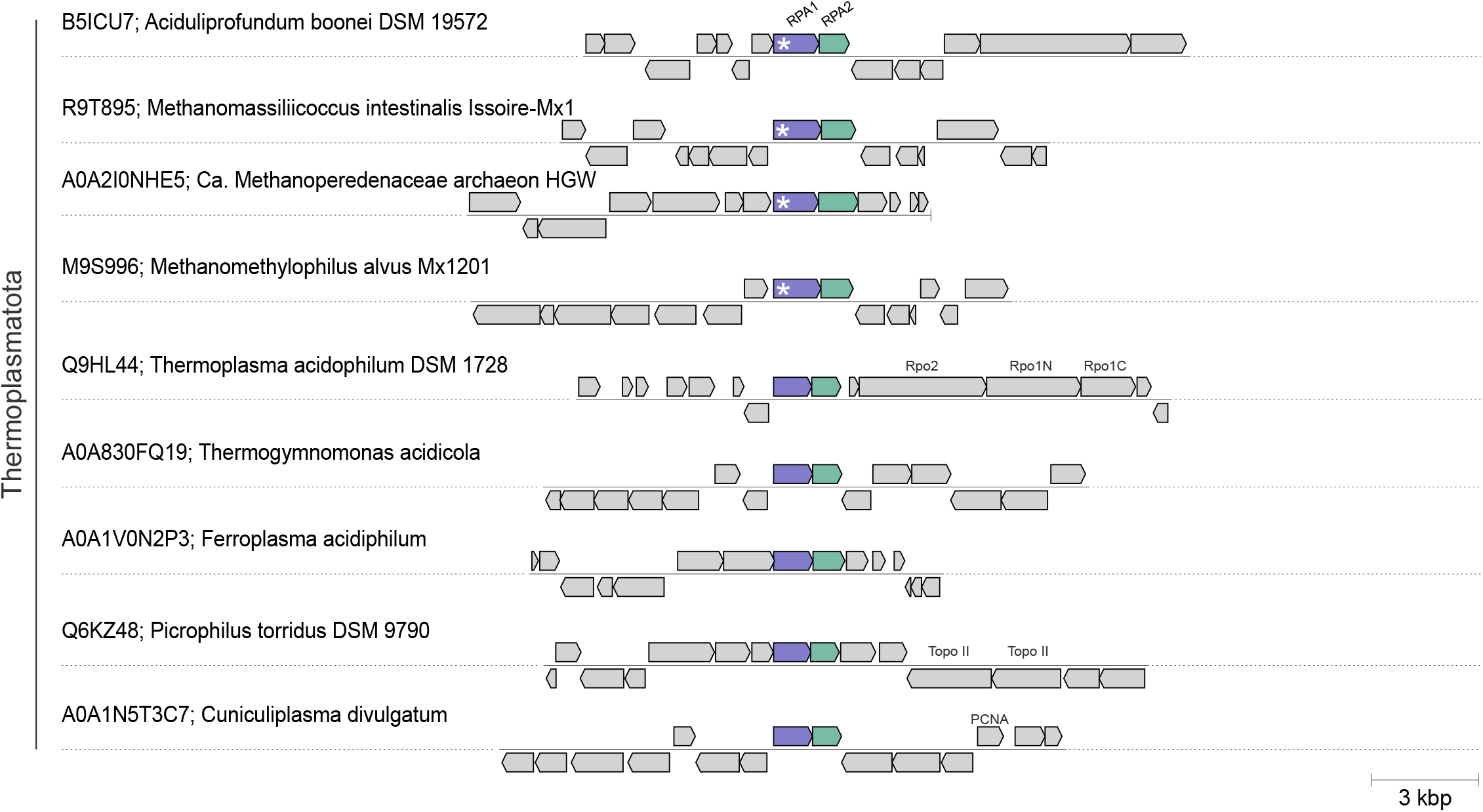
Comparison of genomic loci encoding Rpa1 genes in Archaea. Genomic organizations of the loci encoding Rpa1 genes are shown for representative species from each major archaeal phyla. Each locus is indicated with the UniProt accession number of the Rpa1 gene, followed by the species name. All loci are shown as -7 and +7 genes around Rpa1 gene. Rpa1 genes are embedded within highly variable genomic neighborhoods, which typically vary even for relatively closely related organisms. Presence of the Archaea-specific ARCD is indicated with white asterisks. Strikingly, ARCD is present in all Rpa1 sequences, except for the 5 *Thermoplasmatales* members. It is noteworthy that *Methanomassiliicoccales* within *Thermoplasmatota* do contain ARCD, suggesting that ARCD was lost within *Thermoplasmatales* following their divergence from a common ancestor with *Methanomassiliicoccales*. Interestingly, *Aciduliprofundum boonei* which is even closer to the *Thermoplasmatales* also contain ARCD. DNA metabolism proteins encoding genes are represented in color.

**Figure S4:**
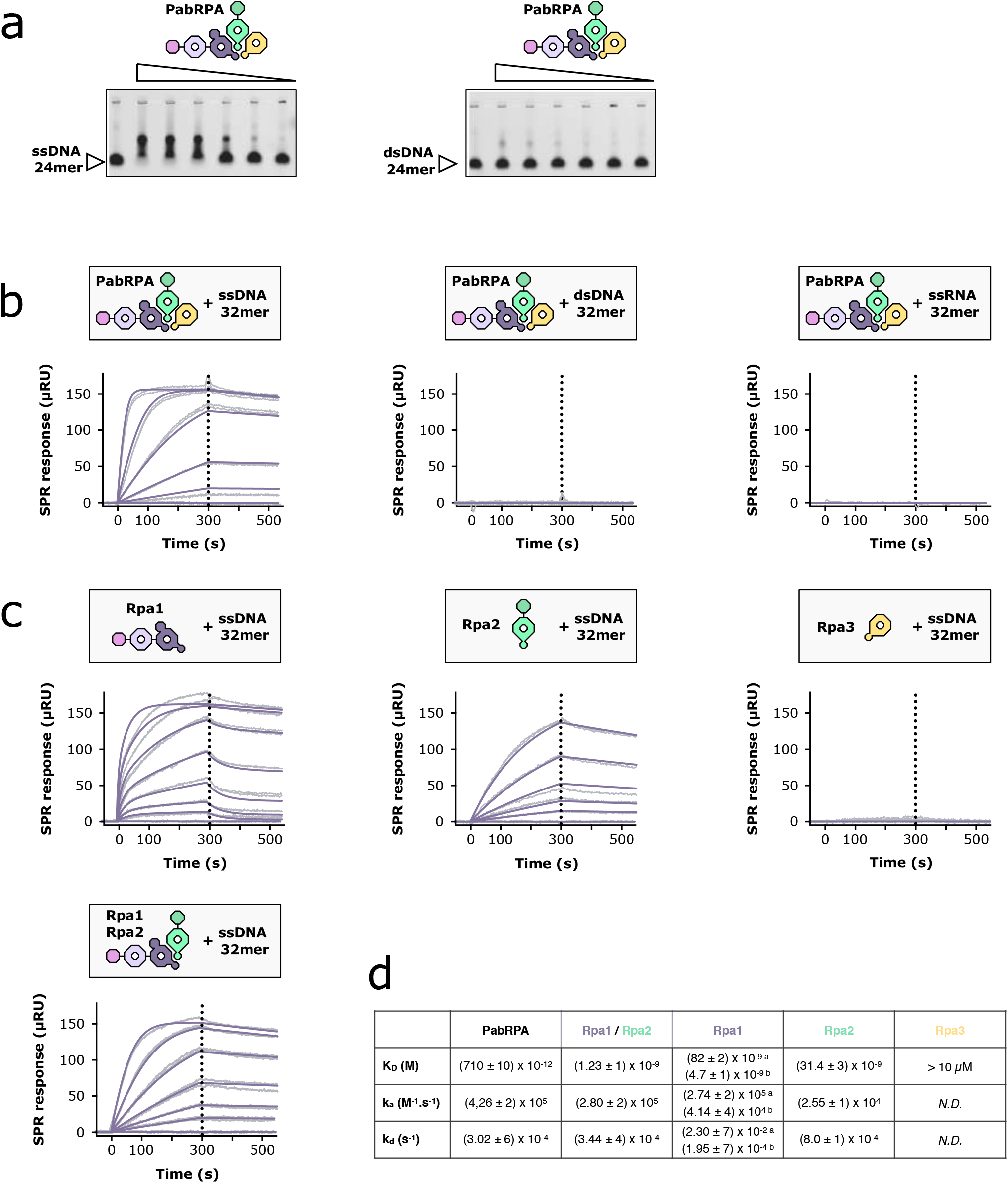
Nucleic-acids binding properties of PabRPA. **a**. Electrophoretic mobility shift assay of ssDNA-24mer and dsDNA-24mer (2.5 pmol) in presence of decreasing concentrations of PabRPA (312, 62, 12, 2.5, 0.5, 0.1 pmol). **b**. PabRPA binding specificity on nucleic acids. Specific binding of PabRPA heterotrimer on immobilized ssDNA-32mer (left), dsDNA-32mer (middle) and ssRNA-32mer measured by surface plasmon resonance (RU: resonance units). **c**. Role of PabRPA subunits on *ss*DNA binding. Specific binding of Rpa1, Rpa2, Rpa3, or Rpa1/Rpa2 complex on immobilized ssDNA-32mer measured by surface plasmon resonance.

**Figure S5:**
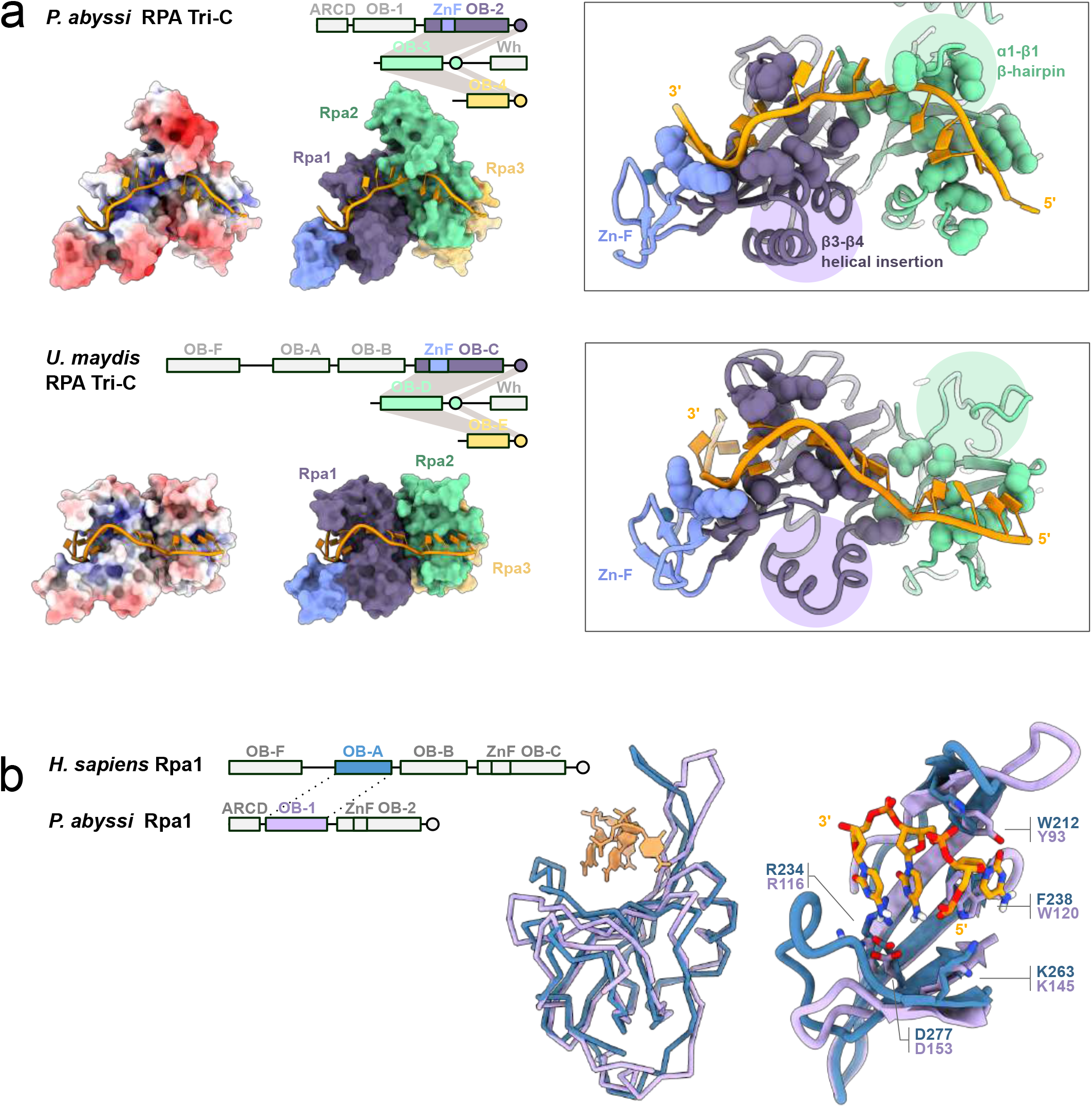
Comparison of archaeal and eukaryotic RPA ssDNA binding determinants. **a**. Comparison of DNA-bound *P. abyssi* and *U. maydis* (4GOP) Tri-C structures with focused views on Rpa1 (purple and blue) and Rpa2 (green). ssDNA-contacting residues are shown as spheres. **b**. Superimposition of the *P. abyssi* Rpa1 OB-1 (in purple) with the *H. sapiens* Rpa1 OB-A (in blue) (1JMC) crystal structures reveals conserved ssDNA-binding amino-acids.

**Figure S6:**
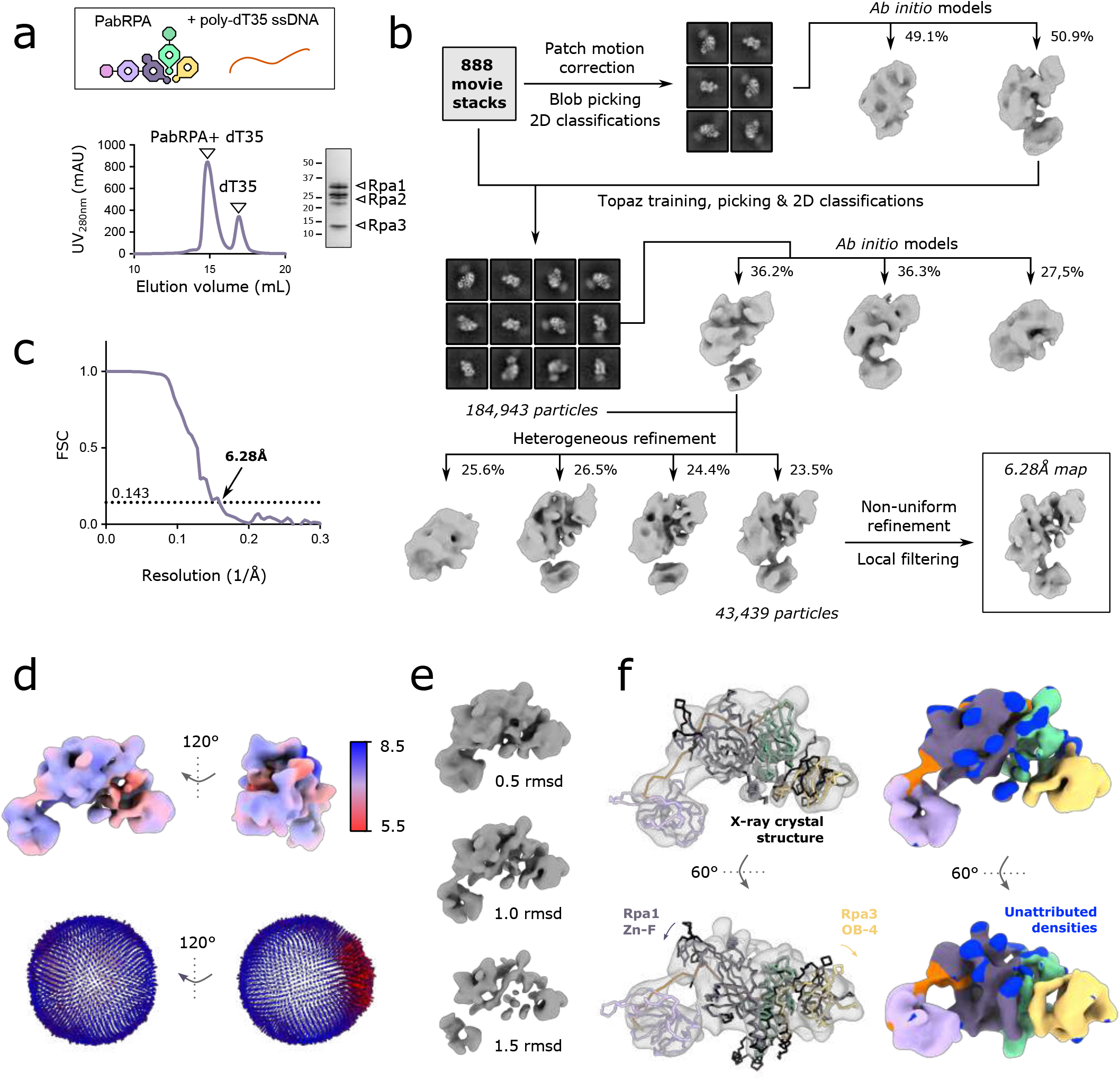
Cryo-EM structure determination of PabRPA bound to a short ssDNA. **a**. Size-exclusion chromatography (SEC) profile (left) and SDS-PAGE analysis (right) of PabRPA bound to a poly-dT35 ssDNA **b**. Data processing workflow (detailed in methods). **c**. Resolution of final reconstructions determined by gold standard FSC at the 0.143 criterion. **d**. Local resolution evaluations of the final reconstructions (top) and Euler angle distributions of particles used for the final reconstructions (bottom). **e**. Cryo-EM maps at varying contour levels. **f**. Comparison of the DNA-bound RPA cryo-EM and crystal structure (left) and cryo-EM map colored by RPA subunits (right).

**Figure S7:**
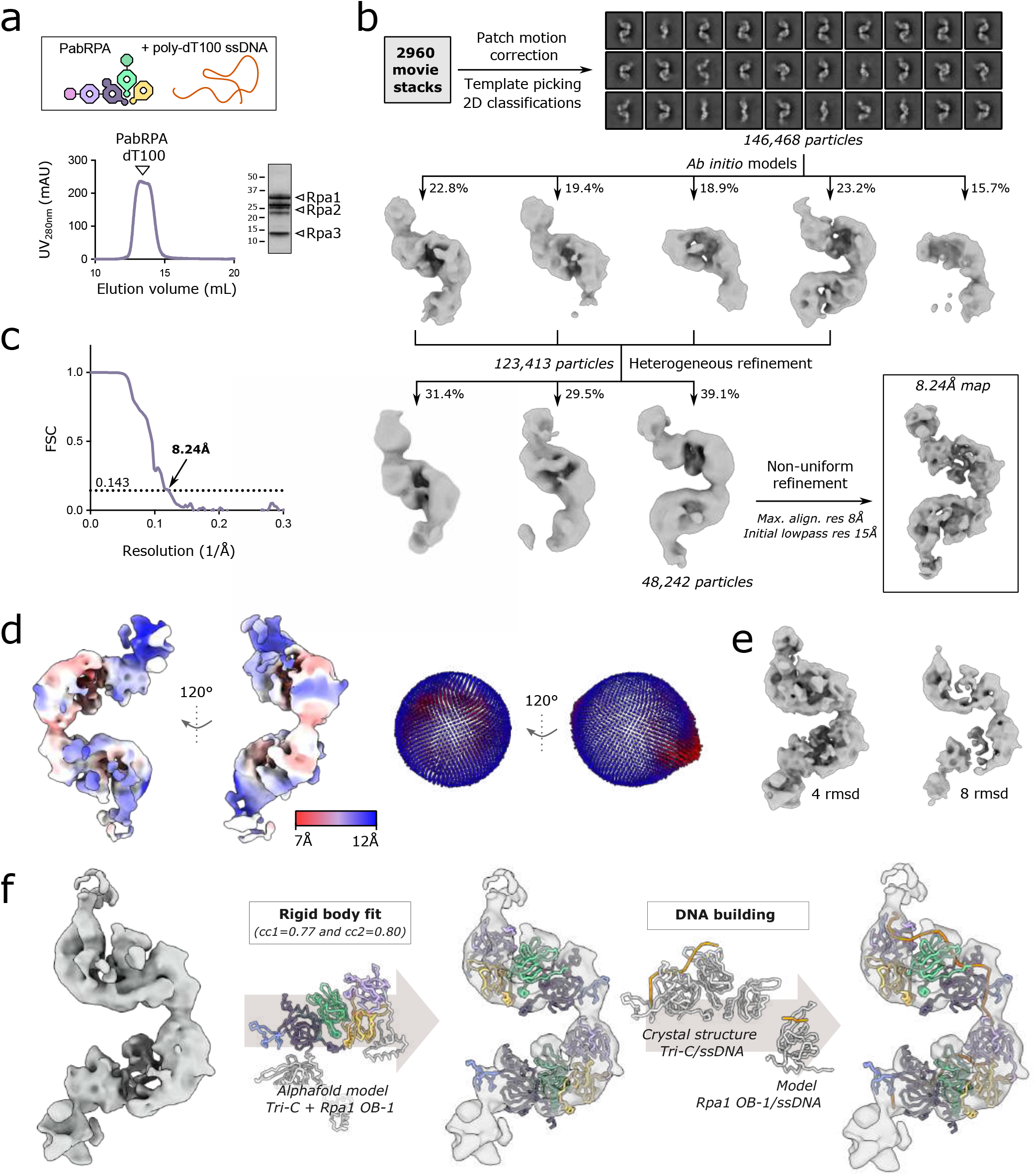
Cryo-EM structure determination of PabRPA bound to a long ssDNA. **a**. Size-exclusion chromatography (SEC) profile (left) and SDS-PAGE analysis (right) of PabRPA bound to a poly-dT100 ssDNA **b**. Data processing workflow (detailed in methods). **c**. Resolution of final reconstructions determined by gold standard FSC at the 0.143 criterion. **d**. Local resolution evaluations of the final reconstructions (left) and Euler angle distributions of particles used for the final reconstructions (right). **e**. Cryo-EM maps at varying contour levels **f**. Model building workflow.

**Figure S8:**
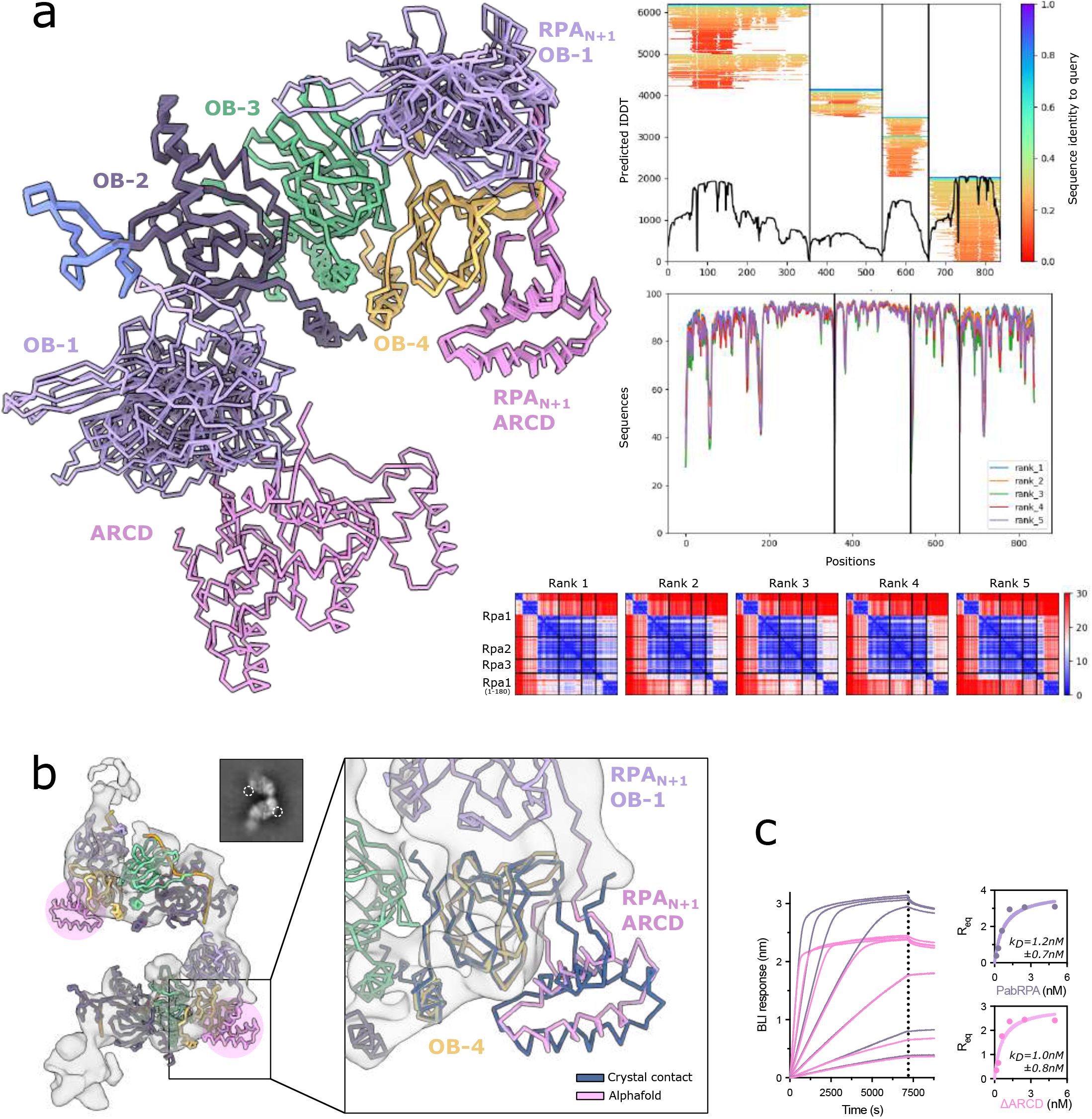
ARCD is required for early-steps of the ssDNA-RPA complex assembly and becomes flexible after binding to ssDNA. **a**. Left: superposition of the five AlphaFold-Multimer predicted models (Evans et al., 2021) for the interaction of one PabRPA-L H heterotrimer with two first domains of PabRPA1 (ARCD-OB1, 1-180). Right: AlphaFold generated plots of MSA sequence coverage (top), of predicted LDDT vs position along the protein sequence (middle) and of predicted aligned error (bottom, where each protein chain of the complexe is plotted against itself and against the other chain). **b**. On left, fit of the RPA(n)/ARCD-OB-1(n+1) AlphaFold model containing the ARCD domain in the cryo-EM map. The ARCD domain is not observed either in the map or in the 2D classes. On right, superimposition with the apo-PabRPA crystal structure. The lattice contact between ARCD and OB-4 fits with the AlphaFold prediction. **c**. Specific binding of PabRPA (in purple) and ΔRCD mutant (in pink) to immobilized poly-dT35 ssDNA measured by biolayer interferometry. Steady-state analysis were performed using the average signal measured at the end of the association steps.

**Figure S9:**
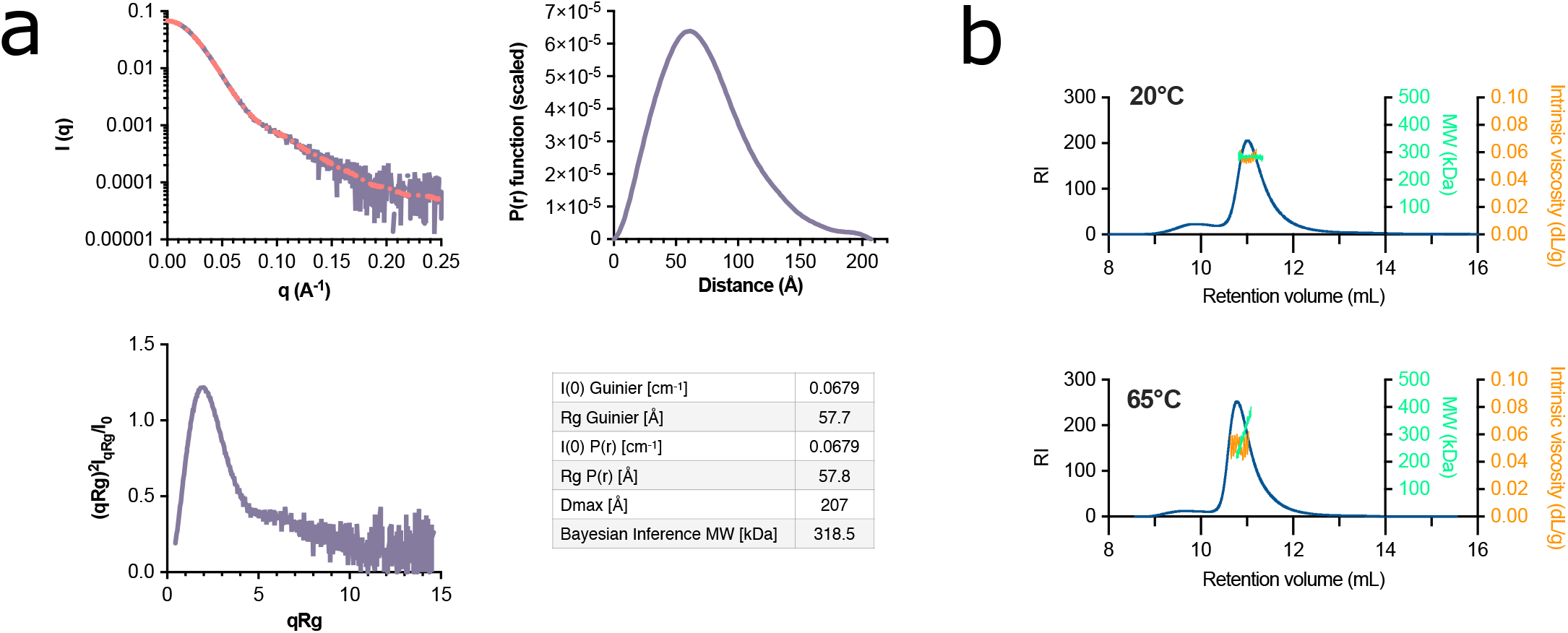
PabRPA oligomerization state in solution. **a**. In solution study of PabRPA structure by SAXS. On top left, SAXS profile of PabRPA (in purple) with superimposition of fitted scattering pattern that allowed the generation of the distance distribution function P(r) (red dotted line). On right, distance distribution functions P(r) derived from the scattering patterns of PabRPA obtained using the program GNOM. On bottom left, dimensionless Kratky plot scattering. Scattering derived parameters are provided in the table. **b**. PabRPA molecular mass (Green) and intrinsic viscosity (orange) measurements by SEC-SLS coupled with an inline viscometer. Measurements were performed at 20°C (higher panel) and 65°C (lower panel).

**Figure S10:**
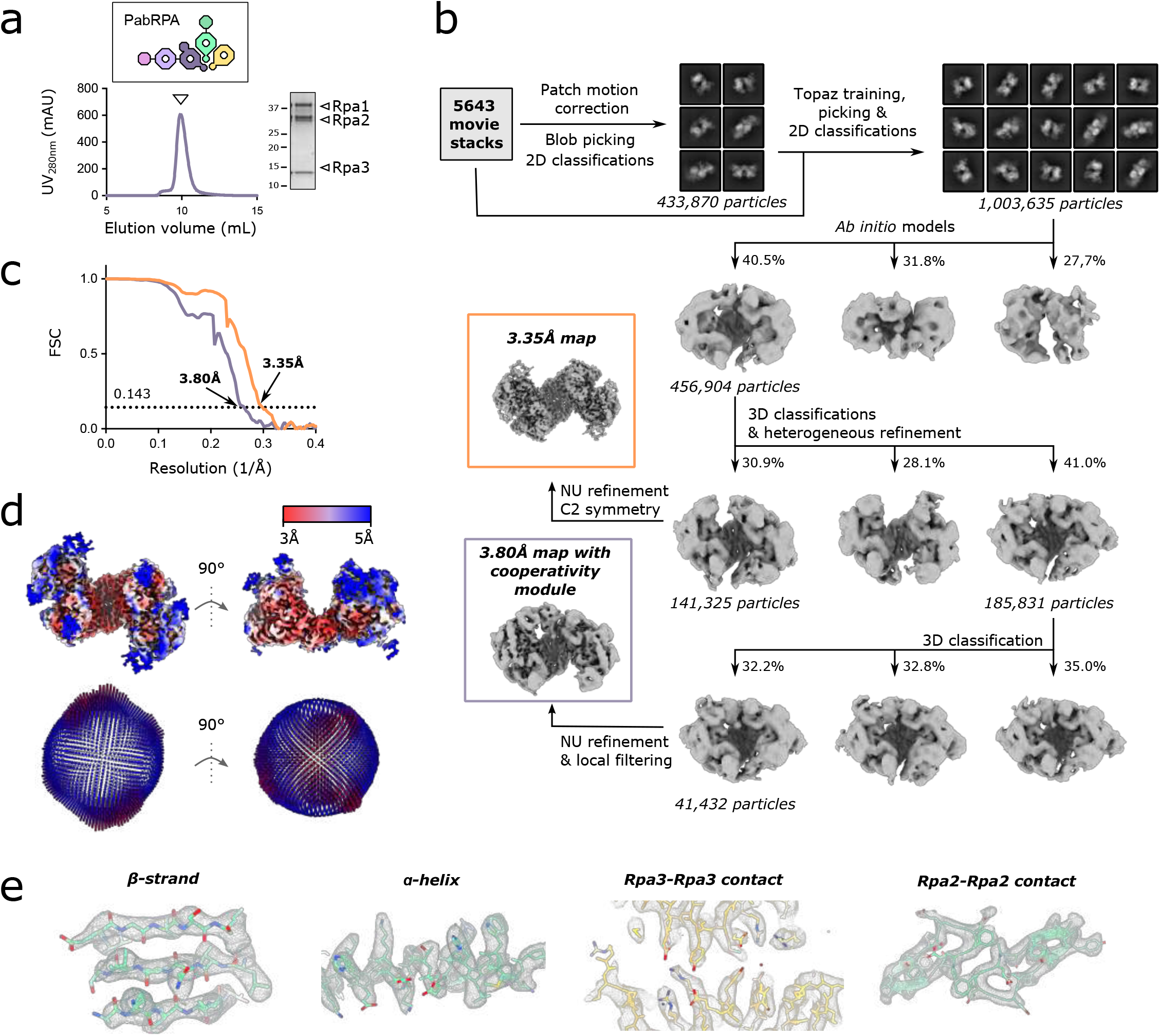
Cryo-EM structure determination of the tetrameric RPA super-structure. **a**. Size-exclusion chromatography (SEC) profile (left) and SDS-PAGE analysis (right) of PabRPA **b**. Data processing workflow (detailed in methods). **c**. Resolution of final reconstructions determined by gold standard FSC at the 0.143 criterion. **d**. Local resolution evaluations of the final reconstructions (top) and Euler angle distributions of particles used for the final reconstructions (bottom). **e**. Representative views of the cryo-EM map showing secondary structure elements and key regions of the tetrameric RPA assembly.

**Table S1:** X-ray diffraction data collection and refinement statistics. Values in parentheses are for highest-resolution shell. Dataset from single crystal used per structure. *Values calculated after truncation by STARANISO. Estimated resolution limits along the three crystallographic directions *a*, b*, c\**.

|  | PabRpa1 cooperativity<br>module<br>(8AA9) | PabRPA<br>(8AAJ) | PabRPA Tri-C/d20T<br>(8AAS) |
| --- | --- | --- | --- |
| **Data collection** |  |  |  |
| Space group | *P* 2<sub>1</sub> | *C* 2 2 2<sub>1</sub> | *P* 3<sub>2</sub> 2 1 |
| Cell dimensions |  |  |  |
| *a*, *b*, *c* (Å) | 52.75, 83.47, 56.19 | 116.53, 169.74, 201.86 | 75.23, 75.23, 215.49 |
| $\alpha$, $\beta$, $\gamma$ (°) | 90, 115.3, 90 | 90, 90, 90 | 90, 90, 120 |
| Wavelength (Å) | 0.97857 | 0.97857 | 0.97856 |
| Resolution (Å) | 50.0 - 1.8 | 49.4 - 3.70 | 48.26 - 3.2 |
|  | (1.85 - 1.8) | (3.80 - 3.70) | (3.28 - 3.2) |
| Estimated resolution limit (Å)* | 1.76, 1.76, 1.91 | 5.46, 4.21, 3.26 | 3.52, 3.52, 3.02 |
| *R*<sub>pim</sub> | 0.025 (0.986) | 0.024 (20.52) | 0.016 (2.752) |
| *R*<sub>merge</sub> | 0.061 (2.43) | 0.123 (89.34) | 0.080 (14.31) |
| *I* / σ*I* | 14.6 (0.8) | 13.0 (0.1) | 22.1 (0.3) |
| CC<sub>1/2</sub> | 0.999 (0.288) | 1.000 (0.130) | 0.999 (0.223) |
| Completeness (%) | 99.9 (99.5) | 99.8 (99.8) | 99.9 (99.9) |
| Redundancy | 7.0 (6.9) | 28.2 (28.4) | 27.1 (27.5) |
| *R*<sub>merge</sub> * | 0.054 (1) | 0.082 (4.31) | 0.067 (4.19) |
| *I* / σ*I* * | 14.6 (1.2) | 21.4 (1.0) | 27.1 (1.0) |
| CC<sub>1/2</sub> * | 0.999 (0.589) | 1.000 (0.542) | 1.000 (0.454) |
| Completeness (%) * | 86.9 (50.5) | 60.1 (13.3) | 80.7 (21.2) |
| **Refinement** |  |  |  |
| Resolution (Å) | 32.25 - 1.80 | 49.4 - 3.70 | 44.84 - 3.2 |
|  | (1.85 - 1.80) | (3.80 - 3.70) | (3.28 - 3.2) |
| No. reflections | 37716 (1857) | 13191 (1036) | 37624 (753) |
| *R*<sub>work</sub> / *R*<sub>free</sub> (%) | 20.86/22.84 | 23.41/25.09 | 26.28/28.90 |
| No. atoms |  |  |  |
| Protein | 2860 | 5304 | 6372 |
| DNA | - | - | - |
| Ligand/ion | 2 | 56 | 258 |
| Water | 290 | - | 335 |
| *B*-factors |  |  |  |
| Protein | 38.6 | 61.9 | 51.04 |
| Ligand/ion | 76.1 | 44.13 | 40.68 |
| Water | 50.8 | 39.49 | 51.69 |
| R.m.s. deviations |  |  |  |
| Bond lengths (Å) | 0.008 | 0.010 | 0.008 |
| Bond angles (°) | 0.95 | 1.45 | 1.18 |

**Table S2:** Cryo-EM data collection, refinement and validation statistics.

|  | PabRPA/dT35<br>(EMDB-15347)<br>(PDB 8ACE) | PabRPA/dT100<br>(EMDB-15350)<br>(PDB 8ACJ) | PabRPA tetramer<br>(EMDB-15300)<br>(PDB 8AAH) | PabRPA tetramer<br>with ARCD-OB-1<br>(EMDB-15301)<br>(PDB 8AAL) |
| --- | --- | --- | --- | --- |
| **Data collection and processing** |  |  |  |  |
| Magnification | 150000 | 150000 | 150000 | 150000 |
| Voltage (kV) | 200 | 200 | 300 | 300 |
| Electron exposure (e<sup>-</sup>/Å<sup>2</sup>) | 40 | 40 | 40 | 40 |
| Defocus range (μm) | -0.8 to -2.8 | -0.8 to -2.8 | -0.8 to -2.8 | -0.8 to -2.8 |
| Pixel size (Å) | 0.96 | 0.96 | 0.86 | 0.86 |
| Symmetry imposed | - | - | C2 | - |
| Initial particle images (no.) | 208094 | 847605 | 1761244 | 1761244 |
| Final particle images (no.) | 43439 | 146468 | 1003635 | 41432 |
| Map resolution (Å) | 6.28 | 8.24 | 3.35 | 3.8 |
| FSC threshold | 0.143 | 0.143 | 0.143 | 0.143 |
| Map resolution range (Å) | 5.5 - 8.5 | 7 - 12 | 3 - 5 | 3 - 5 |
| **Refinement** |  |  |  |  |
| Initial model used (PDB) | 8AA9, 8AAS | 8AA9, 8AAS | 8AAJ | 8AA9, 8AAJ |
| Model resolution (Å) | 8.3 | 10.4 | 3.3 | 3.7 |
| FSC threshold (0.143) |  |  |  |  |
| Map sharpening *B* factor (Å<sup>2</sup>) | 413.2 | 742.8 | 130.0 | 82.4 |
| Model composition |  |  |  |  |
| Non-hydrogen atoms | 4632 | 9428 | 15198 | 16616 |
| Protein residues | 521 | 1048 | 1868 | 2044 |
| Nucleotides | 21 | 46 |  |  |
| Ligands | 1 | 2 | 4 | 4 |
| Mean *B* factors (Å<sup>2</sup>) |  |  |  |  |
| Protein | 777.3 | 620.3 | 45.7 | 278.2 |
| Nucleotides | 810.0 | 586.4 |  |  |
| Ligand | 828.3 | 386.2 | 107.3 | 291.1 |
| R.m.s. deviations |  |  |  |  |
| Bond lengths (Å) | 0.025 | 0.030 | 0.004 | 0.004 |
| Bond angles (°) | 1.945 | 2.196 | 0.685 | 0.697 |
| Validation |  |  |  |  |
| MolProbity score | 2.41 | 2.18 | 1.66 | 1.75 |
| Clashscore | 31.27 | 50.46 | 7.98 | 11.34 |
| Poor rotamers (%) | 1.74 | 0.65 | 0 | 0 |
| Ramachandran plot |  |  |  |  |
| Favored (%) | 96.28 | 98.26 | 96.52 | 96.92 |
| Allowed (%) | 3.33 | 1.36 | 3.26 | 2.8 |
| Disallowed (%) | 0.39 | 0.39 | 0.22 | 0.20 |

**Movie S1: Architecture of PabRPA**

**Movie S2: Structural and functional characterization of PabRPA in complex with ssDNA of different lengths**

**Movie S3: PabRPA forms tetrameric complexes that dissociates upon binding to ssDNA**

## REFERENCES

Adams, P.D., Afonine, P.V., Bunkóczi, G., Chen, V.B., Echols, N., Headd, J.J., Hung, L.-W., Jain, S., Kapral, G.J., Grosse Kunstleve, R.W., McCoy, A.J., Moriarty, N.W., Oeffner, R.D., Read, R.J., Richardson, D.C., Richardson, J.S., Terwilliger, T.C., Zwart, P.H., 2011. The Phenix software for automated determination of macromolecular structures. Methods 55, 94–106. https://doi.org/10.1016/j.ymeth.2011.07.005

Arunkumar, A.I., Stauffer, M.E., Bochkareva, E., Bochkarev, A., Chazin, W.J., 2003. Independent and Coordinated Functions of Replication Protein A Tandem High Affinity Single-stranded DNA Binding Domains. Journal of Biological Chemistry 278, 41077–41082. https://doi.org/10.1074/jbc.M305871200

Ashkenazy, H., Abadi, S., Martz, E., Chay, O., Mayrose, I., Pupko, T., Ben-Tal, N., 2016. ConSurf 2016: an improved methodology to estimate and visualize evolutionary conservation in macromolecules. Nucleic Acids Res 44, W344–W350. https://doi.org/10.1093/nar/gkw408

Bain, F.E., Fischer, L.A., Chen, R., Wold, M.S., 2018. Single-Molecule Analysis of Replication Protein A-DNA Interactions, in: Methods in Enzymology. Elsevier, pp. 439–461. https://doi.org/10.1016/bs.mie.2017.11.016

Barry, E.R., Bell, S.D., 2006. DNA Replication in the Archaea. Microbiology and Molecular Biology Reviews 70, 876–887. https://doi.org/10.1128/MMBR.00029-06

Bepler, T., Morin, A., Rapp, M., Brasch, J., Shapiro, L., Noble, A.J., Berger, B., 2019. Positive-unlabeled convolutional neural networks for particle picking in cryo-electron micrographs. Nature Methods 16, 1153–1160. https://doi.org/10.1038/s41592-019-0575-8

Bianco, P.R., 2017. The tale of SSB. Progress in Biophysics and Molecular Biology 127, 111–118. https://doi.org/10.1016/j.pbiomolbio.2016.11.001

Blackwell, L.J., Borowiec, J.A., 1994. Human replication protein A binds single-stranded DNA in two distinct complexes. Mol Cell Biol 14, 3993–4001. https://doi.org/10.1128/mcb.14.6.3993-4001.1994

Bochkarev, A., Pfuetzner, R.A., Edwards, A.M., Frappier, L., 1997. Structure of the single-stranded-DNA-binding domain of replication protein A bound to DNA. Nature 385, 176–181. https://doi.org/10.1038/385176a0

Bochkareva, E., 2002. Structure of the RPA trimerization core and its role in the multistep DNA-binding mechanism of RPA. The EMBO Journal 21, 1855–1863. https://doi.org/10.1093/emboj/21.7.1855

Bochkareva, E., Frappier, L., Edwards, A.M., Bochkarev, A., 1998. The RPA32 Subunit of Human Replication Protein A Contains a Single-stranded DNA-binding Domain. Journal of Biological Chemistry 273, 3932–3936. https://doi.org/10.1074/jbc.273.7.3932

Bricogne, G., Blanc, E., Brandl, M., Flensburg, C., Keller, P., Paciorek, W., Roversi, P., Sharff, A., Smart, O.S., Vonrhein, C., Womack, T.O., 2017. BUSTER version 2.10.3. Cambridge, United Kingdom: Global Phasing Ltd.

Brill, S.J., Bastin-Shanower, S., 1998. Identification and Characterization of the Fourth Single-Stranded-DNA Binding Domain of Replication Protein A. Mol Cell Biol 18, 7225–7234. https://doi.org/10.1128/MCB.18.12.7225

Brosey, C.A., Chagot, M.-E., Ehrhardt, M., Pretto, D.I., Weiner, B.E., Chazin, W.J., 2009. NMR Analysis of the Architecture and Functional Remodeling of a Modular Multidomain Protein, RPA. J. Am. Chem. Soc. 131, 6346–6347. https://doi.org/10.1021/ja9013634

Brosey, C.A., Soss, S.E., Brooks, S., Yan, C., Ivanov, I., Dorai, K., Chazin, W.J., 2015. Functional Dynamics in Replication Protein A DNA Binding and Protein Recruitment Domains. Structure 23, 1028–1038. https://doi.org/10.1016/j.str.2015.04.008

Brosey, C.A., Yan, C., Tsutakawa, S.E., Heller, W.T., Rambo, R.P., Tainer, J.A., Ivanov, I., Chazin, W.J., 2013. A new structural framework for integrating replication protein A into DNA processing machinery. Nucleic Acids Research 41, 2313–2327. https://doi.org/10.1093/nar/gks1332

Chavas, L.M.G., Gourhant, P., Guimaraes, B.G., Isabet, T., Legrand, P., Lener, R., Montaville, P., Sirigu, S., Thompson, A., 2021. PROXIMA-1 beamline for macromolecular crystallography measurements at Synchrotron SOLEIL. J Synchrotron Rad 28, 970–976. https://doi.org/10.1107/S1600577521002605

Cowtan, K., 2010. Recent developments in classical density modification. Acta Crystallogr D Biol Crystallogr 66, 470–478. https://doi.org/10.1107/S090744490903947X

Deng, X., Habel, J.E., Kabaleeswaran, V., Snell, E.H., Wold, M.S., Borgstahl, G.E.O., 2007. Structure of the Full-length Human RPA14/32 Complex Gives Insights into the Mechanism of DNA Binding and Complex Formation. Journal of Molecular Biology 374, 865–876. https://doi.org/10.1016/j.jmb.2007.09.074

Dueva, R., Iliakis, G., 2020. Replication protein A: a multifunctional protein with roles in DNA replication, repair and beyond. NAR Cancer 2, zcaa022. https://doi.org/10.1093/narcan/zcaa022

Duggin, I.G., Bell, S.D., 2006. The Chromosome Replication Machinery of the Archaeon Sulfolobus solfataricus. Journal of Biological Chemistry 281, 15029–15032. https://doi.org/10.1074/jbc.R500029200

Durand, D., Vivès, C., Cannella, D., Pérez, J., Pebay-Peyroula, E., Vachette, P., Fieschi, F., 2010. NADPH oxidase activator p67phox behaves in solution as a multidomain protein with semi-flexible linkers. Journal of Structural Biology 169, 45–53. https://doi.org/10.1016/j.jsb.2009.08.009

Dutta, A., Stillman, B., 1992. cdc2 family kinases phosphorylate a human cell DNA replication factor, RPA, and activate DNA replication. The EMBO Journal 11, 2189–2199. https://doi.org/10.1002/j.1460-2075.1992.tb05278.x

Emsley, P., Cowtan, K., 2004. Coot : model-building tools for molecular graphics. Acta Crystallogr D Biol Crystallogr 60, 2126–2132. https://doi.org/10.1107/S0907444904019158

Emsley, P., Lohkamp, B., Scott, W.G., Cowtan, K., 2010. Features and development of Coot. Acta Crystallogr D Biol Crystallogr 66, 486–501. https://doi.org/10.1107/S0907444910007493

Evans, R., O’Neill, M., Pritzel, A., Antropova, N., Senior, A., Green, T., Zidek, A., Bates, R., Blackwell, S., Yim, J., Ronneberger, O., Bodenstein, S., Zielinski, M., Bridgland, A., Potapenko, A., Cowie, A., Tunyasuvunakool, K., Jain, R., Clancy, E., Kohli, P., Jumper, J., Hassabis, D., 2021. Protein complex prediction with AlphaFold-Multimer (preprint). Bioinformatics. https://doi.org/10.1101/2021.10.04.463034

Fan, J., Pavletich, N.P., 2012. Structure and conformational change of a replication protein A heterotrimer bound to ssDNA. Genes & Development 26, 2337–2347. https://doi.org/10.1101/gad.194787.112

Feldkamp, M.D., Frank, A.O., Kennedy, J.P., Patrone, J.D., Vangamudi, B., Waterson, A.G., Fesik, S.W., Chazin, W.J., 2013. Surface Reengineering of RPA70N Enables Cocrystallization with an Inhibitor of the Replication Protein A Interaction Motif of ATR Interacting Protein. Biochemistry 52, 6515–6524. https://doi.org/10.1021/bi400542z

Gamsjaeger, R., Kariawasam, R., Gimenez, A.X., Touma, C., McIlwain, E., Bernardo, R.E., Shepherd, N.E., Ataide, S.F., Dong, Q., Richard, D.J., White, M.F., Cubeddu, L., 2015. The structural basis of DNA binding by the single-stranded DNA-binding protein from Sulfolobus solfataricus. Biochemical Journal 465, 337–346. https://doi.org/10.1042/BJ20141140

Holm, L., 2020. DALI and the persistence of protein shape. Protein Science 29, 128–140. https://doi.org/10.1002/pro.3749

Jiang, X., Klimovich, V., Arunkumar, A.I., Hysinger, E.B., Wang, Y., Ott, R.D., Guler, G.D., Weiner, B., Chazin, W.J., Fanning, E., 2006. Structural mechanism of RPA loading on DNA during activation of a simple pre-replication complex. EMBO J 25, 5516–5526. https://doi.org/10.1038/sj.emboj.7601432

Jumper, J., Evans, R., Pritzel, A., Green, T., Figurnov, M., Ronneberger, O., Tunyasuvunakool, K., Bates, R., Zidek, A., Potapenko, A., Bridgland, A., Meyer, C., Kohl, S.A.A., Ballard, A.J., Cowie, A., Romera-Paredes, B., Nikolov, S., Jain, R., Adler, J., Back, T., Petersen, S., Reiman, D., Clancy, E., Zielinski, M., Steinegger, M., Pacholska, M., Berghammer, T., Bodenstein, S., Silver, D., Vinyals, O., Senior, A.W., Kavukcuoglu, K., Kohli, P., Hassabis, D., 2021. Highly accurate protein structure prediction with AlphaFold. Nature 596, 583–589. https://doi.org/10.1038/s41586-021-03819-2

Jurrus, E., Engel, D., Star, K., Monson, K., Brandi, J., Felberg, L.E., Brookes, D.H., Wilson, L., Chen, J., Liles, K., Chun, M., Li, P., Gohara, D.W., Dolinsky, T., Konecny, R., Koes, D.R., Nielsen, J.E., Head-Gordon, T., Geng, W., Krasny, R., Wei, G., Holst, M.J., McCammon, J.A., Baker, N.A., 2018. Improvements to the APBS biomolecular solvation software suite. Protein Science 27, 112–128. https://doi.org/10.1002/pro.3280

Kabsch, W., 2010. XDS. Acta Crystallogr D Biol Crystallogr 66, 125–132. https://doi.org/10.1107/S0907444909047337

Kerr, I.D., 2003. Insights into ssDNA recognition by the OB fold from a structural and thermodynamic study of Sulfolobus SSB protein. The EMBO Journal 22, 2561–2570. https://doi.org/10.1093/emboj/cdg272

Kõivomägi, M., Valk, E., Venta, R., Iofik, A., Lepiku, M., Balog, E.R.M., Rubin, S.M., Morgan, D.O., Loog, M., 2011. Cascades of multisite phosphorylation control Sic1 destruction at the onset of S phase. Nature 480, 128–131. https://doi.org/10.1038/nature10560

Komori, K., Ishino, Y., 2001. Replication Protein A in Pyrococcus furiosus Is Involved in Homologous DNA Recombination. Journal of Biological Chemistry 276, 25654–25660. https://doi.org/10.1074/jbc.M102423200

Krissinel, E., Henrick, K., 2007. Inference of Macromolecular Assemblies from Crystalline State. Journal of Molecular Biology 372, 774–797. https://doi.org/10.1016/j.jmb.2007.05.022

Legrand, P., 2017. XDSME: XDS Made Easier. GitHub (https://github.com/legrandp/xdsme).

Lim, C.J., Barbour, A.T., Zaug, A.J., Goodrich, K.J., McKay, A.E., Wuttke, D.S., Cech, T.R., 2020. The structure of human CST reveals a decameric assembly bound to telomeric DNA. Science 368, 1081–1085. https://doi.org/10.1126/science.aaz9649

Liu, Y., Makarova, K.S., Huang, W.-C., Wolf, Y.I., Nikolskaya, A.N., Zhang, X., Cai, M., Zhang, C.-J., Xu, W., Luo, Z., Cheng, L., Koonin, E.V., Li, M., 2021. Expanded diversity of Asgard archaea and their relationships with eukaryotes. Nature 593, 553–557. https://doi.org/10.1038/s41586-021-03494-3

MacNeill, S.A., 2021. Remote Homology Detection Identifies a Eukaryotic RPA DBD-C-like DNA Binding Domain as a Conserved Feature of Archaeal Rpa1-Like Proteins. Frontiers in Molecular Biosciences 8. https://doi.org/10.3389/fmolb.2021.675229

Makarova, K.S., Koonin, E.V., 2013. Archaeology of Eukaryotic DNA Replication. Cold Spring Harbor Perspectives in Biology 5, a012963–a012963. https://doi.org/10.1101/cshperspect.a012963

Marceau, A.H., 2012. Functions of Single-Strand DNA-Binding Proteins in DNA Replication, Recombination, and Repair, in: Keck, J.L. (Ed.), Single-Stranded DNA Binding Proteins. Humana Press, Totowa, NJ, pp. 1–21. https://doi.org/10.1007/978-1-62703-032-8_1

Maréchal, A., Zou, L., 2015. RPA-coated single-stranded DNA as a platform for post-translational modifications in the DNA damage response. Cell Res 25, 9–23. https://doi.org/10.1038/cr.2014.147

McCoy, A.J., Grosse-Kunstleve, R.W., Adams, P.D., Winn, M.D., Storoni, L.C., Read, R.J., 2007. Phaser crystallographic software. J Appl Crystallogr 40, 658–674. https://doi.org/10.1107/S0021889807021206

Meyer, R.R., Laine, P.S., 1990. The single-stranded DNA-binding protein of Escherichia coli. Microbiol Rev 54, 342–380. https://doi.org/10.1128/mr.54.4.342-380.1990

Mirdita, M., Schütze, K., Moriwaki, Y., Heo, L., Ovchinnikov, S., Steinegger, M., 2021. ColabFold - Making protein folding accessible to all (preprint). Bioinformatics. https://doi.org/10.1101/2021.08.15.456425

Mistry, J., Chuguransky, S., Williams, L., Qureshi, M., Salazar, G.A., Sonnhammer, E.L.L., Tosatto, S.C.E., Paladin, L., Raj, S., Richardson, L.J., Finn, R.D., Bateman, A., 2021. Pfam: The protein families database in 2021. Nucleic Acids Research 49, D412–D419. https://doi.org/10.1093/nar/gkaa913

Miyake, Y., Nakamura, M., Nabetani, A., Shimamura, S., Tamura, M., Yonehara, S., Saito, M., Ishikawa, F., 2009. RPA-like Mammalian Ctc1-Stn1-Ten1 Complex Binds to Single-Stranded DNA and Protects Telomeres Independently of the Pot1 Pathway. Molecular Cell 36, 193–206. https://doi.org/10.1016/j.molcel.2009.08.009

Morten, M.J., Peregrina, J.R., Figueira-Gonzalez, M., Ackermann, K., Bode, B.E., White, M.F., Penedo, J.C., 2015. Binding dynamics of a monomeric SSB protein to DNA: a single-molecule multi-process approach. Nucleic Acids Res 43, 10907–10924. https://doi.org/10.1093/nar/gkv1225

Nagata, M., Ishino, S., Yamagami, T., Ishino, Y., 2019. Replication protein A complex in Thermococcus kodakarensis interacts with DNA polymerases and helps their effective strand synthesis. Bioscience, Biotechnology, and Biochemistry 83, 695–704. https://doi.org/10.1080/09168451.2018.1559722

Oliveira, M.T. (Ed.), 2021. Single Stranded DNA Binding Proteins, Methods in Molecular Biology. Springer US, New York, NY. https://doi.org/10.1007/978-1-0716-1290-3

Pan, Z.Q., Amin, A.A., Gibbs, E., Niu, H., Hurwitz, J., 1994. Phosphorylation of the p34 subunit of human single-stranded-DNA-binding protein in cyclin A-activated G1 extracts is catalyzed by cdk-cyclin A complex and DNA-dependent protein kinase. Proc. Natl. Acad. Sci. U.S.A. 91, 8343–8347. https://doi.org/10.1073/pnas.91.18.8343

Park, C.-J., 2005. Solution structure of the DNA-binding domain of RPA from Saccharomyces cerevisiae and its interaction with single-stranded DNA and SV40 T antigen. Nucleic Acids Research 33, 4172–4181. https://doi.org/10.1093/nar/gki736

Pérez, J., Vachette, P., Russo, D., Desmadril, M., Durand, D., 2001. Heat-induced unfolding of neocarzinostatin, a small all-B protein investigated by small-angle X-ray scattering 1 1Edited by M. F. Moody. Journal of Molecular Biology 308, 721–743. https://doi.org/10.1006/jmbi.2001.4611

Pettersen, E.F., Goddard, T.D., Huang, C.C., Meng, E.C., Couch, G.S., Croll, T.I., Morris, J.H., Ferrin, T.E., 2021. UCSF CHIMERAX : Structure visualization for researchers, educators, and developers. Protein Science 30, 70–82. https://doi.org/10.1002/pro.3943

Punjani, A., Rubinstein, J.L., Fleet, D.J., Brubaker, M.A., 2017. cryoSPARC: algorithms for rapid unsupervised cryo-EM structure determination. Nat Methods 14, 290–296. https://doi.org/10.1038/nmeth.4169

Raymann, K., Forterre, P., Brochier-Armanet, C., Gribaldo, S., 2014. Global Phylogenomic Analysis Disentangles the Complex Evolutionary History of DNA Replication in Archaea. Genome Biology and Evolution 6, 192–212. https://doi.org/10.1093/gbe/evu004

Robbins, J.B., Murphy, M.C., White, B.A., Mackie, R.I., Ha, T., Cann, I.K.O., 2004. Functional Analysis of Multiple Single-stranded DNA-binding Proteins from Methanosarcina acetivorans and Their Effects on DNA Synthesis by DNA Polymerase BI. Journal of Biological Chemistry 279, 6315–6326. https://doi.org/10.1074/jbc.M304491200

Seeber, A., Hegnauer, A.M., Hustedt, N., Deshpande, I., Poli, J., Eglinger, J., Pasero, P., Gut, H., Shinohara, M., Hopfner, K.-P., Shimada, K., Gasser, S.M., 2016. RPA Mediates Recruitment of MRX to Forks and Double-Strand Breaks to Hold Sister Chromatids Together. Molecular Cell 64, 951–966. https://doi.org/10.1016/j.molcel.2016.10.032

Sheldrick, G.M., 2008. A short history of SHELX. Acta Crystallogr A Found Crystallogr 64, 112–122. https://doi.org/10.1107/S0108767307043930

Sibenaller, Z.A., Sorensen, B.R., Wold, M.S., 1998. The 32-and 14-Kilodalton Subunits of Replication Protein A Are Responsible for Species-Specific Interactions with Single-Stranded DNA. Biochemistry 37, 12496–12506. https://doi.org/10.1021/bi981110+

Spang, A., Saw, J.H., Jørgensen, S.L., Zaremba-Niedzwiedzka, K., Martijn, J., Lind, A.E., van Eijk, R., Schleper, C., Guy, L., Ettema, T.J.G., 2015. Complex archaea that bridge the gap between prokaryotes and eukaryotes. Nature 521, 173–179. https://doi.org/10.1038/nature14447

Stephan, H., Concannon, C., Kremmer, E., Carty, M.P., Nasheuer, H.-P., 2009. Ionizing radiation-dependent and independent phosphorylation of the 32-kDa subunit of replication protein A during mitosis. Nucleic Acids Research 37, 6028–6041. https://doi.org/10.1093/nar/gkp605

Stroud, A., Liddell, S., Allers, T., 2012. Genetic and Biochemical Identification of a Novel Single-Stranded DNA-Binding Complex in Haloferax volcanii. Frontiers in Microbiology 3. https://doi.org/10.3389/fmicb.2012.00224

Surovtseva, Y.V., Churikov, D., Boltz, K.A., Song, X., Lamb, J.C., Warrington, R., Leehy, K., Heacock, M., Price, C.M., Shippen, D.E., 2009. Conserved Telomere Maintenance Component 1 Interacts with STN1 and Maintains Chromosome Ends in Higher Eukaryotes. Molecular Cell 36, 207–218. https://doi.org/10.1016/j.molcel.2009.09.017

Svergun, D.I., 1992. Determination of the regularization parameter in indirect-transform methods using perceptual criteria. J Appl Crystallogr 25, 495–503. https://doi.org/10.1107/S0021889892001663

Taib, N., Gribaldo, S., MacNeill, S.A., 2021. Single-Stranded DNA-Binding Proteins in the Archaea, in: Oliveira, M.T. (Ed.), Single Stranded DNA Binding Proteins. Springer US, New York, NY, pp. 23–47. https://doi.org/10.1007/978-1-0716-1290-3_2

Terwilliger, T.C., Poon, B.K., Afonine, P.V., Schlicksup, C.J., Croll, T.I., Millán, C., Richardson, J.S., Read, R.J., Adams, P.D., 2022. Improved AlphaFold modeling with implicit experimental information (preprint). Biochemistry. https://doi.org/10.1101/2022.01.07.475350

Theobald, D.L., Mitton-Fry, R.M., Wuttke, D.S., 2003. Nucleic Acid Recognition by OB-Fold Proteins. Annu. Rev. Biophys. Biomol. Struct. 32, 115–133. https://doi.org/10.1146/annurev.biophys.32.110601.142506

Tickle, I.J., Flensburg, C., Keller, P., Paciorek, W., Sharff, A., Vonrhein, C., Bricogne, G., 2018. STARANISO (http://staraniso.globalphasing.org/cgi-bin/staraniso.cgi). Cambridge, United Kingdom: Global Phasing Ltd.

Vagin, A., Teplyakov, A., 1997. MOLREP : an Automated Program for Molecular Replacement. Journal of Applied Crystallography 30, 1022–1025. https://doi.org/10.1107/S0021889897006766

Wadsworth, R.I.M., 2001. Identification and properties of the crenarchaeal single-stranded DNA binding protein from Sulfolobus solfataricus. Nucleic Acids Research 29, 914–920. https://doi.org/10.1093/nar/29.4.914

Wold, M.S., 1997. REPLICATION PROTEIN A: A Heterotrimeric, Single-Stranded DNA-Binding Protein Required for Eukaryotic DNA Metabolism. Annual Review of Biochemistry 66, 61–92. https://doi.org/10.1146/annurev.biochem.66.1.61

Yates, L.A., Aramayo, R.J., Pokhrel, N., Caldwell, C.C., Kaplan, J.A., Perera, R.L., Spies, M., Antony, E., Zhang, X., 2018. A structural and dynamic model for the assembly of Replication Protein A on single-stranded DNA. Nat Commun 9, 5447. https://doi.org/10.1038/s41467-018-07883-7

